# Single cell approaches define the murine leptomeninges:cortical brain interface as a distinct cellular neighborhood comprised of neural and nonneural cell types

**DOI:** 10.1101/2025.01.24.634752

**Authors:** Sarah N. Ebert, Christine Eisner, Konstantina Karamboulas, Louis-Philippe Bernier, David R. Kaplan, Brian A. MacVicar, Freda D. Miller

## Abstract

The interface between the brain surface and the adjacent meninges is a selective barrier regulating fluid, protein and immune cell exchange between the CNS and periphery. However, the cell types that form this important interface are not yet fully defined. To address this limitation, we have used single cell RNA-sequencing (scRNA-seq) and single cell spatial transcriptomics together with morphological lineage tracing and immunostaining to analyze the adult murine cortex. We show that the cortical interface is comprised of three major cell types, leptomeningeal cells, border astrocytes and tissue-resident macrophages. On the peripheral side the interface is comprised of transcriptionally-distinct PDGFRα-positive leptomeningeal mesenchymal cells that are intermingled with macrophages. This leptomeningeal pial layer is lined by a population of transcriptionally-distinct border astrocytes. The interface neighborhood is rich in growth factor mRNAs, including many leptomeningeal ligands predicted to act on both the border astrocytes and macrophages. On the CNS side of the interface is the relatively cell-sparse cortical layer one containing interneurons, microglia, parenchymal astrocytes, oligodendrocyte precursor cells and oligodendrocytes. Except for the border astrocytes, layer one cells are not closely-associated with the interface, suggesting that secreted ligands may be the major way the brain interface communicates with the underlying cortical parenchyma. Thus, our data provide a molecular/cellular resource describing the brain interface cell types and their interactions, thereby enabling future studies asking how this distinct cellular compartment regulates CNS:periphery interactions.

**Significance Statement:** Recent years have seen significant progress in identifying the diverse cell types within the meningeal space. However, the mechanisms by which these cells interact with glial and neuronal cells in layer one of the adult murine cortex remain poorly understood. During development, communication between radial precursors and meningeal layers is crucial for proper brain formation, but the role of this interaction in adulthood is still unclear. Additionally, how resident immune cells in the leptomeningeal space signal to layer one cortical cells or meningeal mesenchymal cells during homeostasis remains an open question. Understanding the identity, location, and interactions of these cells is essential for unraveling the complex dynamics at this critical brain interface.

## Introduction

The interface between the brain surface and the periphery ensures maintenance of a distinct neural environment by acting as a selective barrier regulating the passage of fluid, proteins, and inflammatory cells. On the superficial side of this interface are the meninges, a complex tissue comprised of blood vessels, nerves, immune cells, and several distinct populations of mesenchymal cells (Betsholtz et al., 2024; Como et al., 2023; Rua and McGavern, 2018). The superficial-most meningeal layer is the dura, which is adjacent to the skull and functions as a connective tissue stroma (Kolabas et al., 2023; Rustenhoven et al., 2021). Closer to the brain are the leptomeninges, which are comprised of arachnoid and arachnoid barrier cells, and, immediately adjacent to the brain, pial cells (Betsholtz et al., 2024; Como et al., 2023; DeSisto et al., 2020). In humans, these distinct layers have long been characterized anatomically, but we have only recently begun to understand the meninges at the molecular/cellular level, largely as a consequence of single cell transcriptional studies (Kearns et al., 2023; Wang et al., 2022). Nonetheless, we still don’t understand how leptomeningeal cells interact with CNS cells to form the brain:periphery interface nor do we fully understand what happens when this interface is broached following traumatic injury, viral and bacterial attack, neurodegenerative disorders, or neurosurgery (Aydin et al., 2023; Derk et al., 2022; Ewing-Crystal et al., 2024; Roth et al., 2014; Turtzo et al., 2020).

What then do we know about the brain interface cells and how might we obtain a better understanding of their characteristics? We know that the adult brain interface includes leptomeningeal pial cells on the peripheral side and astrocytes on the CNS side. One idea is that this astrocyte:pial cell interaction may be conceptually analogous to the much better-studied astrocyte:vasculature interaction. Indeed, several recent studies demonstrated that leptomeningeal cells are associated with large blood vessels that penetrate the brain from the outside (Bonney et al., 2022; Jones et al., 2023; Pietilä et al., 2023). However, there is little direct evidence that this astrocyte:pial cell interaction participates in blood-brain barrier function. Instead, several studies indicate that pial cells function as part of a brain surface signaling center, at least during embryogenesis when they provide a basal anchor for cortical radial precursors and secrete ligands that regulate multiple aspects of cortical development (Choe et al., 2014; Como et al., 2023; Dasgupta and Jeong, 2019; Hartmann et al., 1998; Izen et al., 2018; Sievers et al., 1994; Zelco et al., 2021). Whether adult pial cells play similar roles is unclear, although some studies suggest that they might. For example, when cortical leptomeningeal cells were removed locally this caused perturbations both in cortical layer one and in cortical midline structures suggesting that leptomeningeal cells may play multiple roles in the adult brain (Nikouei et al., 2024).

These studies highlight the importance of acquiring a molecular/cellular understanding of adult brain interface cells and their interactions. Here, we have addressed this key issue by creating a single cell RNA-sequencing and spatial transcriptomic resource focused on the cortical brain interface and adjacent cortical layer one. Our data define three distinct cell types within the brain interface, leptomeningeal mesenchymal cells, border astrocytes that are transcriptionally-distinct from parenchymal astrocytes, and a population of tissue-resident border macrophages. We show that these three cell types comprise a tightly-organized cellular neighborhood, and that this neighborhood is not closely-associated with any layer one cortical cell type other than the border astrocytes. Moreover, we identify the ligands and ligand receptors expressed by the three interface cell types and use these to predict paracrine interactions within the interface neighborhood. Thus, our data provide a molecular/cellular resource describing the brain interface cell types and their potential interactions, thereby enabling future studies asking about the functions of this distinct cellular compartment during homeostasis and following injury.

## Materials and Methods

### Animals

All animal protocols were approved by the Animal Care Committees of the relevant institutions in accordance with national animal care policies. Animals had free access to rodent chow and water and were housed in a temperature and humidity-controlled environment on a 12-hour light-dark cycle. All mice were healthy with no obvious behavioral phenotypes. For all studies, adult (8–16 week old) mice of either sex were used and mice were randomly allocated to experimental groups. Wild type C57BL6 mice were purchased from Charles River Laboratories (strain codes: 027) or Jackson Laboratories (Strain Code: 000664). *PdgfraCreERT* (B6 N.Cg-Tg(Pdgfra-cre/ERT)467Dbe/J; JAX stock #018280) RRID:IMSR_JAX:018280 (Kang et al., 2010), *R26-LSL-TdTomato* (B6.Cg-Gt(ROSA)26Sortm9(CAG-tdTomato)Hze/J; JAX stock #007909) RRID:IMSR_JAX:007909 (Madisen et al., 2010), *Tmem119CreERT2* (C57BL/6-Tmem119em1(cre/ERT2)Gfng/J;, JAX stock #031820) RRID:IMSR_JAX:031820 (Kaiser and Feng, 2019), were obtained from Jackson Laboratories. All mice were bred and genotyped as recommended by Jackson Laboratories.

For experiments involving inducible Cre recombinase mouse strains, tamoxifen (Sigma) was dissolved in a sunflower oil/ethanol mixture (9:1) at 30 mg/mL. Mice were injected intraperitoneally with 3 mg/day for five consecutive days, followed by a 10 day wash out period and then collected for imaging.

### Tissue preparation, immunostaining and microscopy

For immunostaining analyses mice were perfused trans-cardially with ice cold 1x PBS and then with 4% PFA. The head, lower jaw, and skin were removed from the skull, and the snout removed just rostral to the eyes. Excess muscle tissue was also removed from the cranium. Finally, the ventral portion of the upper palate was removed exposing the ventral surface of the brain. The skull and brain were post-fixed in 4% PFA for 18-24 hours at 4°C, and then washed in PBS. To decalcify the skull, the tissue was incubated in 0.5M EDTA (pH 8.0) at 4°C for 4 days, or until bone was sufficiently soft. Tissue was then incubated in 15% Sucrose for 48 hours at 4°C, followed by an incubation in 30% sucrose for 48 hours or until tissue had sunk. Tissue was then incubated in OCT for 4 hours at 4°C with slow rotation and snap fixed in fresh OCT. Brains were sectioned sagittally and tissue was collected from the M1 region line preserving the orientation of skull, meninges, and brain. Sections were 12-14 uM thick and stored at −80°C until staining.

For immunostaining, sections were dried for 15 minutes at 37°C or until all condensation disappeared, incubated in 1X PBS for 10 minutes to rehydrate, and then blocked using 5% BSA (Jackson ImmunoResearch) containing 0.3% Triton X-100 (Fisher) in PBS for 1 h at room temperature. Sections were incubated with primary antibodies diluted in 5% BSA in PBS overnight at 4°C. Sections were washed 3 times with PBS prior to application of appropriate secondary antibodies at optimized concentrations (see Antibodies section). Following a 1-hour room temperature incubation in the dark, sections were washed 3 times with PBS, counterstained with DAPI (1:1000) for 5 minutes at room temperature, and washed a further 3 times with PBS. Coverslips were applied to tissue with Vectashield and sealed.

Images were collected with either a Zeiss Axio Imager M2 system with an X-Cite 120 LED light source, Apotome 3 and C11440 Hamamatsu camera or a Zeiss LSM900 with an Airyscan 2 detector (for representative images). Images were processed using Fiji (Schindelin et al., 2012) (RRID: SCR_002285). Gamma adjustments for GFAP immunostaining in Figure 3 were used to brighten processes against very bright cell bodies (0.75). Similarly for *Tmem119CreERT2*:Tdtomato strain imaging in the same Figure, the Tdt channel was also adjusted (0.75).

### Single cell isolation and scRNA-seq

For single cell isolation of meningeal cells, tissue was isolated from the inside of the skull (bone associated) and the cortical surface of the brain (cortical surface associated). For the former, the top of the skull above the cortex with associated tissue was removed from the brain surface and the meningeal tissue was carefully removed from the inner side of the skull under a microscope in ice-cold PBS. The tissue was chopped using a scalpel and collected into a 15mL falcon tube containing 2mL of Collagenase D (1.5U/mL) (Roche ref 11088882001)/Dispase II (2.4U/ml) (Roche ref 4942078001) + 10ul/ml CaCl2 (stock 250mM), and incubated at 37°C with rotation/agitation for 45 minutes or until digested. Every 15 minutes the sample was checked and vortexed. Post digest, the cells were triturated with a p200 followed by a polished glass pipette, topped up to 15mL volume with HBSS + 2mM EDTA. It was then filtered through a 70 uM filter, spun down 7 minutes at 800g, supernatant removed, and resuspended in 1mL 0.2% BSA in HBSS for FACS sorting and sequencing.

For the cortical surface associated digest, the top half of the dorsal cortex was glued onto a 1cm diameter circular coverslip in a 24 well plate, rinsed with HBSS, the HBSS was then replaced with 1-1.5mL of collagenase dispase solution as above. The plate was incubated at 37°C with rotation/agitation for 30-45 minutes or until digested. Every 15 minutes the sample was checked, and the well solution was ‘washed’ over the brain 5 times with a pipette for each sample. Once digestion was complete, the supernatant from each well was combined into a 15mL falcon tube, triturated with a p200 and polished glass pipette, and topped up to 15mL with HBSS + 2mM EDTA. The sample was filtered through a 70 uM filter, spun down for 7 minutes at 800g, supernatant removed, and resuspended in 1mL 0.2% BSA in HBSS for FACS sorting and sequencing.

For all samples, viable cells were incubated in DRAQ5 (1:3000) and propidium iodide (PI) (1:1000) and sorted by flow cytometry. Cells that were DRAQ5-positive and PI-negative were collected in DMEM, L-Glutamine, and B27 and submitted for single cell isolation and processing using the 10X Genomics Chromium system as per the manufacturer’s guidelines. The resultant libraries were sequenced on an Illumina NextSeq500. FASTQ sequencing reads were processed, aligned to wild type mouse genome (mm10) and converted to digital gene expression matrices using the Cell Ranger count function within the Cell Ranger Single-Cell Software Suite (RRID: SCR_017344) with settings as recommended by the manufacturer (https://support.10xgenomics.com/single-cell-gene-expression/software/overview/welcome) (Zheng et al., 2017). Two bone associated and four cortical surface-associated samples were sequenced that each included tissue from equal numbers of mice of each sex. The two bone-associated runs included tissue from 10 mice and 16 mice total. The four cortical surface-associated runs included tissue from 10, 16, 18 and 18 mice total.

### scRNA-seq analysis

Dataset count matrices were processed using a previously published pipeline (Borrett et al., 2020; Innes and Bader, 2019; Willis et al., 2025). In brief, datasets were filtered to remove low expressing cells, and cells with a high mitochondrial gene expression profile. Using R Project for Statistical Computing (Version 4.1,2)((R Core Team, 2021), Datasets were normalized using scran (version 1.28.0), and data was transferred into Seurat (version 4.0.1, RRID:SCR_016341) (Hao et al., 2021; Lun et al., 2016). PCA analysis was performed using highly variable genes and UMAPs generated using the top principal components detected in the datasets. Once clustered, all datasets were subsetted to remove clusters with low transcript counts and the raw count matrices for the selected cells were run back through the pipeline omitting filtering steps. Cell cycle scores were computed as part of the pipeline using Cyclone, part of the scran package. Two-dimensional t-SNE projections were generated using the top principal components (RunTSNE, Seurat). The same principal components were also used to execute SNN-Cliq-inspired clustering (FindClusters Seurat) with iteratively increasing resolution until the number of differentially expressed genes (FindMarkers Seurat, where p < 0.01 family wise error rate (FWER) Holm method) between the most similar clusters reached approximately 30 genes. All individual or merged datasets were analyzed with the most conservative resolution selected based on clustering that best aligned with the expression of well-defined markers for each cell type.

We used gene expression overlays on Uniform Manifold Approximation and Projection (UMAP) and annotated clusters and cell type identities based on the following markers; *Pdgfra, Sox10, Cspg4, Enpp6* for OPCs, *Mbp, Mog, Mag, Sox10* for oligodendrocytes, *Pdgfrb, Myh11, Mylk, Acta2* for blood vessel-associated mesenchymal cells (VAMCs), *Pdgfrb, Rgs5, Cspg4, Kcnj8* for pericytes, *Ptprc, Cd68, Cd44* for all immune cells, *Lyv1, Mrc1, Dab2, Clec10a* for border macrophages, *Clec4n, Folr1, Ccr2, Lgals3* for dural macrophages, *Cxcr2* for neutrophils, *Cd19* for B cells, *Nkg7, Cd3e, Cd4, Cd8a* for T cells, *Aif1, Sall1, Tmem119, Cx3cr1, Aif1, Iba1*for microglia, *Aqp4, Slc1a3, Arxes2, Cxcl14* for astrocytes, *Myoc, Gfap* for border astrocytes, *Plp1, Sox2, Ngfr, Sox10, Sema3d* for Schwann cells, *Pecam1, Plvap, Ly6c1, Cdh5* for endothelial cells, *Ttr, Foxj1* for choroid plexus, *Pdgfra, Col1a1, Gjb6, Gjb2, Bmp7, Bmp6, Wnt5a, Lama1, Ptgds* for leptomeningeal mesenchymal cells, *Pdgfra, Col1a1, H19, Matn4, Clec11a, Shisa3* for dural mesenchymal cells, *Gad1, Gad2* for interneurons, and *Slc17a7, Slc17a6* for excitatory neurons.

### Pearson correlation analysis

Pearson correlation of scRNAseq data was preformed between leptomeningeal and dural mesenchymal cells by averaging the expression of each gene across all cells, and then determining the Pearson correlation coefficient with the cor.test function in R.

### Dataset merger and batch correction

To generate the merged cortical surface associated datasets and the bone associated datasets, all cells passing library thresholds were extracted from each dataset and the raw transcriptomes were run through the pipeline as indicated above. Following PCA, batch correction was used for the cortical surface associated datasets using one iteration of Harmony to correct for sequencing differences (Korsunsky et al. 2019). No batch correction was needed for bone associated datasets. When subsetting *Pdgfra*-positive clusters to generate the merged meningeal mesenchymal cell dataset, leptomeningeal and dural meningeal mesenchymal cells were extracted and merged, and raw transcriptomes run through the pipeline as indicated above. One iteration of Harmony was performed to correct for the aforementioned sequencing differences within the cortical surface associated datasets.

For astrocyte datasets, a similar process was used for each of the 6 previously-published datasets (GEO GSE148611; Hasel et al., 2021). These were merged with previously-published P60 Grey Matter and P60 Cortex datasets (GEO GSE255405; Dennis et al., 2024) along with astrocytes from cortical surface datasets described in this manuscript. One iteration of Harmony was performed to correct for sequencing depth differences between datasets. Batch ID was assigned with all 6 Hasel datasets under one batch, and each subsequent dataset with its own Batch ID.

For the leptomeningeal mesenchymal, dural mesenchymal, and border astrocyte cell merge dataset, as well as the leptomeningeal mesenchymal, border astrocyte, and border macrophage merge dataset, appropriate clusters were extracted from the datasets of origin, merged together as described above, and one iteration of Harmony was performed to correct for sequencing differences.

### Ligand-receptor analysis

Ligand-receptor analysis was performed essentially as described in Willis et al. (2025) using a curated ligand-receptor database (Toma et al., 2020) and the CCINX package (Isserlin et al., 2020). Ligands and receptors were included when they were detectably expressed in >5% of the relevant cell types. Cytoscape (v3.9.1, SCR_003032) was used to visualize the predictive ligand-receptor communication models, where ligands included in the model are presented in a central panel of nodes and edges connecting them to their source and target cell type (Shannon et al., 2003).

### Xenium in situ based single cell spatial transcriptomics

Single cell multiplexed in situ gene expression analysis was performed using the Xenium platform as described in (Ma et al., 2024; Willis et al., 2025). Analysis was performed on adult C57Bl/6 mouse brain tissue with or without an accompanying dural dissection. Two C57Bl/6 adult mice, one male and one female, were used with the Adult Brain Panel with custom add on (WHMTAZ). Four adult mice (2 male and 2 female) with an included dural dissection (described below), were used with the Adult Mesenchymal Custom Panel (9XXV3X).

Following RNAse-free removal of brains, fresh tissue was flash frozen in OCT embedding matrix and stored at −70°C. Subsequently, 10 µm cryosections were mounted onto Xenium slides (chemistry v2) following 10X Genomics guidelines. All tissues were equilibrated to −21°C in a cryostat prior to sectioning. Each coronal section was collected within the M1 cortical region, rostral to the hippocampus. One section was collected from each of the two adult mice used with the adult brain panel, two sections were collected from each of the male and female mice that were used with the mesenchymal panel (Janesick et al., 2023). For slides prepared for the mesenchymal probeset, the dura was carefully dissected off the skull, on top of the brain, prior to embedding. Slides were stored at −70°C prior to subsequent preparation steps.

Sectioned tissue was processed according to the Xenium workflow protocol for fresh frozen tissue. To fix and permeabilize the tissue, sections were incubated for 1 minute at 37°C, then fixed in 4% PFA (Electron Microscopy Sciences) for 30 minutes at room temperature. Sections were washed in RNAse-free 1X PBS for one minute (Thermo Fisher), and subjected to a series of permeabilization steps and PBS washes. Permeabilization steps included 1% SDS incubation for 2 minutes and a chilled 70% methanol incubation for 1 hour. Following permeabilization, slides were washed twice with PBS and then placed in PBS/0.05% Tween-20 (PBS-T, Thermo Fisher). Sections were hybridized with either a ‘Brain Panel’ probe solution containing 347 genes (see Extended Table 9), or a ‘Mesenchymal Panel’ with 491 genes (see Extended Table 9) in TE-buffer, for 22 hours at 50°C. Afterwards, sections underwent several PBS washes and were incubated at 37°C in post-hybridization wash buffer. Following a series of PBS-T washes, sections were incubated in Xenium ligation enzymes for 2 hours at 37°C. Following multiple rounds of PBS-T washes, probe amplification was achieved by incubating sections for 2 hours at 30°C in Xenium amplification enzyme solution. Sections were washed twice in TE buffer, and stored overnight at 4°C. Autofluorescence quenching and nuclei staining were next performed. Sections were washed in PBS, incubated in reducing agent for 10 minutes, washed in 70% and 100% ethanol, and incubated in Xenium autofluorescence solution for 10 minutes. Sections were then washed three times in 100% ethanol, dried at 37°C for 5 minutes and rehydrated via a series of PBS and PBS-T washes. Nuclear staining buffer was added to sections for 1 minute and sections washed 4 times in PBS-T. Samples were loaded into the Xenium analyzer instrument (v1.7.6.0) and subjected to several cycles of reagent application, probe hybridization, imaging, and probe removal. Pre-processing of captured Z-stack images was performed using the Xenium on-board analysis pipeline. In brief, a custom Xenium codebook (described below) was used to decode imaged fluorescent puncta into transcripts, with each decoded fluorescence signature assigned a codeword which was associated to a target gene. Quality scores (Q-Scores) were assigned to transcripts based on maximum likelihood codewords compared to the likelihood of other sub-optimal codewords, to provide the confidence in each decoded transcripts assigned identity. Only transcripts with a Q-score ≥ 20 were included in downstream analysis. Negative control codewords (codewords that do not correspond to any probe) and negative control probes (probes included in the panel which do not match any biological sequence) were utilized to assess decoding accuracy and assay specificity, respectively. On-instrument cell segmentation was performed based on 3D DAPI morphology. Nuclei identification and cell boundaries were subsequently flattened into a 2D mask, cell IDs allocated to each identified cell, and transcripts assigned to cell IDs based on their X-Y co-ordinates. The boundary for expansion around the nucleus was set for 2 um manually for each section to account for the tight packing of cells at the brain interface.

### Single cell spatial transcriptomic data analysis

Standardized Xenium output files were exported for downstream analyses. In addition, 3 datasets from Willis et al. 2025 (GEO GSE266689) that were run using the same 347 gene brain probeset were downloaded and analyzed in the same way. Data were visualized in Xenium Explorer (version 3.0, RRID: SCR_025847), with regions of interest (ROIs) defined using the freehand select tool and cell IDs exported (Janesick et al. 2023). ROIs were drawn to include the brain surface, and the cortex down to the bottom of layer 2. Transcript count, feature, coordinate and cell ID data were imported to R and Seurat objects created (v5.0.1, RRID:SCR_016341) via the LoadXenium function (Hao et al., 2024). Subsets were then produced from object data using the previously exported cell IDs to include only cells contained within our ROI, and these regions were concatenated where multiple ROIs were specified on one section. All downstream Xenium analysis was conducted in R package Seurat. Initial quality checks assessed cells based on the number of detected genes and total transcript counts per cell. Low quality cells with greater than +/-2.5 standard deviations from the mean in either of these parameters were excluded.

Filtered data were normalized using SCTransform and PCA based on highly variable genes to capture main sources of variation within the data and reduce dimensionality. To construct a nearest neighbor (SNN) graph, Seurat Find Neighbors was used with principal components identified through PCA. A Louvain clustering algorithm based in Seurat (Find Clusters) was applied to partition cells into distinct clusters based on transcriptomic profile. Further analysis was performed on the smallest, yet still biologically meaningful clustering resolution.

UMAPs and image dimensional reduction plots visualizing cell co-ordinates (ImageDimPlot) were employed to visualize the data. Cell clusters were annotated using well defined marker genes for specific cell types. For slides analyzed with the brain probeset, leptomeningeal cells were identified with *Lama1, Slc22a6, Gjb2, Dpp4*, astrocytes with *Aqp4, Eva1*, and border astrocytes specifically with *Gfap, Myoc*. Microglia were identified with *Tmem119, Cx3cr1, Cd68*, and border macrophages with *Cybb*, *Cd53.* Excitatory neurons were identified with *Dpyd, Rspo2, Adamts2, Cdh13* and various populations of inhibitory neurons with *Gad1, Gad2, Lamp5, Pvalb, Vip and Sst.* OPCs and oligodendrocytes were identified with *Olig1, Olig2, Sox10*, and OPCs specifically with *Cspg4, Pdgfra.* Vascular associated mesenchymal cells were identified with *Acta2, Pdgfrb*, and pericytes with *Cspg4, Pdgfrb*. Endothelial cells were identified with *Pecam1* and *Cdh5*.

For the mesenchymal probeset, leptomeningeal cells were identified with *Pdgfra, Lama1, Slc22a6, Gjb2, Dpp4* and dural mesenchymal cells with *Pdgfra, H19, Matn4, Shisa3*, and *Clec11a*. Astrocytes were identified with *Aqp4, Agt, Arxes2*, and border astrocytes specifically with *Gfap, Myoc*. Microglia were identified with *Tmem119, Cx3cr1, Cd68*, and border macrophages with *Mrc1, Lyz2.* Excitatory neurons were identified with *Plppr3, Rab40b, Smad3, Bdnf* and inhibitory neurons with *Dlx5, Cryab, Prox1* (upper interneurons) and *Dlx5, Mkx, Galnt9* (lower interneuron). OPCs and oligodendrocytes were identified with *Erbb3, Sox10*, and OPCs specifically with *Cspg4, Pdgfra.* Vascular associated mesenchymal cells were identified with *Myh11, Pdgfrb*, and pericytes with *Kcnj8, Rgs5, Pdgfrb*.

Each Xenium field of view (FOV) was processed and annotated individually, and Seurat SelectIntegrationFeatures, FindIntegrationAnchors and IntegrateData functions used to merge FOVs. After merging, Normalization, PCA, SNN analysis, clustering, and annotation were performed on merged datasets and/or subsets as described above, resulting in one merged Seurat object for each panel. The mesenchymal probeset included datasets from 6 sections/4 independent mice total, and the brain probeset analysis included 2 sections/2 independent mice plus 3 sections/3 independent mice from Willis et al. 2025 (GEO GSE266689) (5 sections/5 mice total).

### Cellular proximity analysis

To perform nearest neighbor analysis we used the Giotto toolkit as described by (Dries et al., 2021). We created spatial networks using both Delauney and kNN methods with a maximum distance of 400, and used the kNN method with the *cellProximityEnrichment* function to create a cell proximities frequency table. Each cell was assigned a label such as astrocyte or leptomeninges, and we used a permutation test to compare our interaction (cell proximity count) data to a null distribution created from a random permutation of cell labels while keeping the positions fixed. Hence, the overall number of cells and proportion of each cell type are preserved but the spatial arrangement is randomized. We used 1000 simulations to create the null distribution of expected cell proximity counts for comparison with our observed data. We used the fdr adjustment method for our p value adjustment. P_higher (enrichment) and p_lower (depletion) (raw one sided p values) assessed the probability that the observed frequency is either significantly higher (enrichment) or lower (depletion). This is shown further in the Fruchterman layout using the *cellProximityNetwork* function, where green lines indicate a depleted interaction when compared to null distribution, and black indicate an enriched interaction.

### Antibodies

We used the following primary antibodies for immunostaining; goat anti-PDGFRα (R&D Systems Cat# AF1062; RRID: AB_2236897), rabbit anti-Laminin (Abcam, Cat. Ab11575; RRID: AB_298179), rabbit anti-Cx30 (Invitrogen, Cat. 71-2000; RRID AB_2533979), rabbit anti-Cx26 (Invitrogen, Cat. 51-2800 Lot#XA340183; RRID AB_2533903), rat anti-GFAP (Invitrogen, Cat. 13-0300, Lot#: WI331291; AB_2532994), rabbit anti-MATN (LSBio, Cat. LS-C108492-0.1, Lot#217995 RRID: RRID:AB_10637203). Fluorescently labeled highly cross-absorbed secondary antibodies were purchased from Jackson ImmunoResearch and used at a dilution of 1:300 - 1:500. Donkey Anti-Rat Alexa Flour 488 (Cat#: 712-545-153; RRID: AB_2340684), Donkey Anti-Rabbit Alexa Flour 555 (Cat#: 711-565-152; RRID: AB_3095471), Donkey Anti-Goat Alexa Flour 647 (Cat#: 705-605-147, RRID: AB_2340437).

### Experimental Design and Statistical Tests

PCA analysis was performed using highly variable genes and UMAPs generated using the top principal components detected in the datasets. Two-dimensional t-SNE projections were generated using the top principal components (RunTSNE, Seurat). The same principal components were also used to execute SNN-Cliq-inspired clustering (FindClusters Seurat) with iteratively increasing resolution until the number of differentially expressed genes (FindMarkers Seurat, where p < 0.01 family wise error rate (FWER), Holm method) between the most similar clusters reached approximately 30 genes. We used the Pearson correlation coefficient test (cor.test function in R) to determine similarities between leptomeningeal and dural mesenchymal cells. For the cell proximity analysis, we used permutation test to compare a generated random null distribution to our collected data.

## Results

### scRNA-seq identifies cell types within the meningeal:brain interface of the adult murine cerebral cortex

To ask about cells that comprise the cortical brain surface interface, we initially performed single cell RNA-sequencing (scRNA-seq) and used two different dissection protocols to obtain a global overview of the cells at this interface (see Materials and Methods). In one approach, we isolated cortical surface-associated cells, including the leptomeninges, by dissecting the top of the cortex, adhering the ventral surface to a coverslip, and then liberating single cells enzymatically. In the second approach, we enriched for bone-associated cells such as the dura by removing the skull, dissecting the associated connective tissue, and enzymatically dissociating single cells. For both dissections, we used FACs to select for viable cells, and then isolated and sequenced single cells using the 10X Genomics Chromium platform. In total, we performed 4 separate runs for the cortical surface and 2 runs for bone-associated cells, using 5-10 mice per run.

The resultant transcriptomes were analyzed using a previously described pipeline (see Materials and Methods for details). After filtering the datasets for low quality cells with few expressed genes or high mitochondrial proportions and cell doublets, we obtained 810 to 2709 single cell transcriptomes in each of the six independent datasets (9436 cells total). We then merged the transcriptomes from the bone-associated datasets and those from the cortical surface datasets separately. We used genes with high variance to compute principal components as inputs for projecting cells in two-dimensions using UMAPs and clustering performed using a shared nearest neighbors-cliq (SNN-cliq)-inspired approach built into the Seurat R package at a range of resolutions (Hao et al., 2024). Since there was some segregation between the four different cortical surface datasets, we normalized the merged dataset using one iteration of Harmony batch correction (Korsunsky et al., 2019) (Fig. 1-1A). We then used gene expression overlays for well-validated marker genes to define cell types.

**Figure 1.**
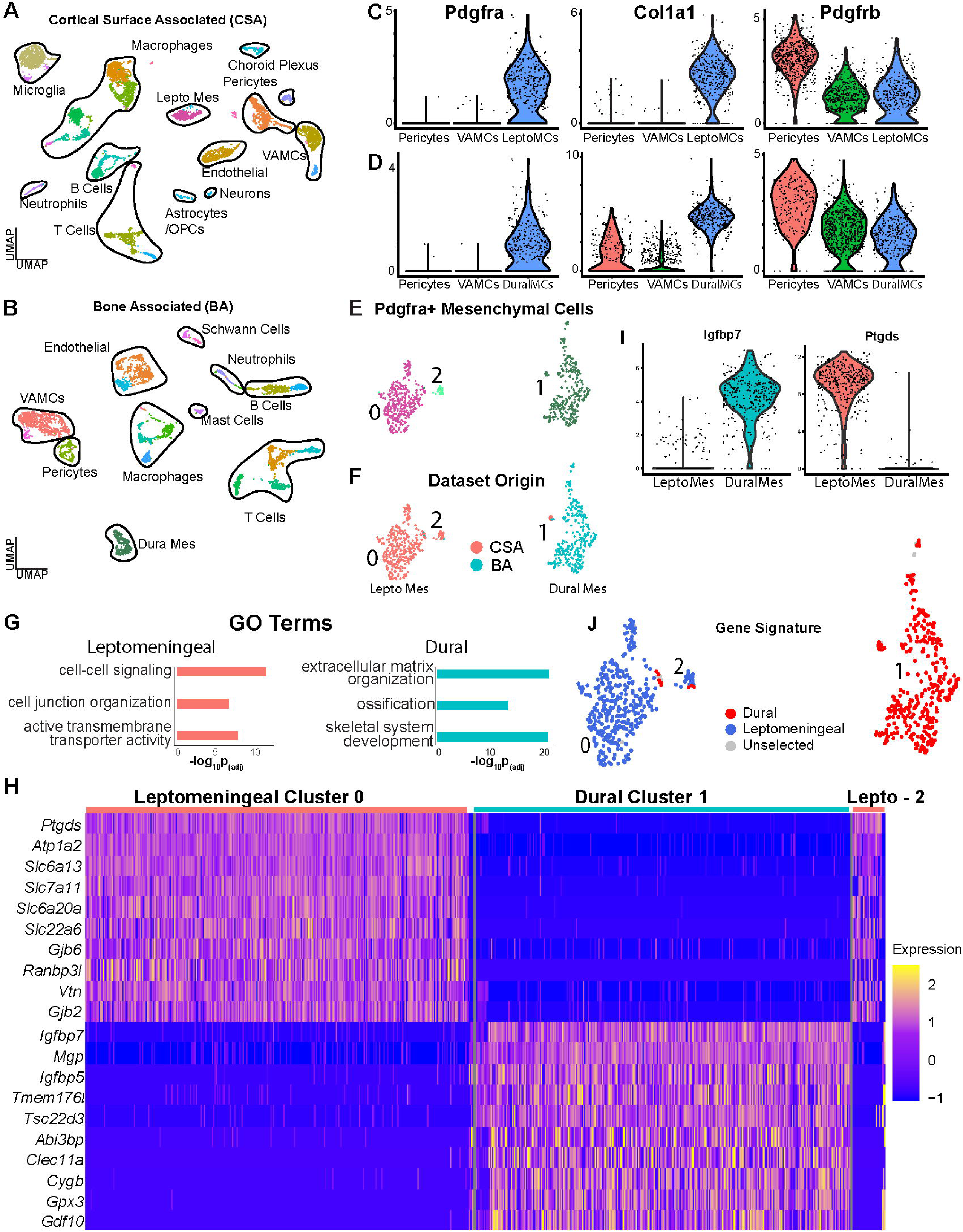
*scRNA-seq identifies distinct populations of cortical brain interface versus bone-associated cells*. Also see Extended Figure 1-1 and Extended Tables 1 and 2. **(A, B)** UMAP visualization of all transcriptomes identified in the cortical surface-associated (CSA) (A) or cortical bone-associated (BA) (B) datasets as analyzed by scRNA-seq. Shown are the mergers of 4 independent surface-associated runs (A) and 2 independent bone-associated runs (B). The merged CSA dataet in (A) was batch-corrected using 1 iteration of Harmony. Transcriptionally-distinct clusters are colour-coded, and plots are annotated for cell types corresponding to these clusters, as determined by analysis of well-characterized marker genes. VAMCs = vasculature-associated mesenchymal cells, Lepto Mes = leptomeningeal mesenchymal cells, Dura Mes = dural mesenchymal cells. **(C, D)** Violin plots showing expression of *Pdgfra, Pdgfrb* and *Col1a1* in leptomeningeal (C; colored blue) and dural (D; colored blue) mesenchymal cells relative to the *Pdgfra*-negative pericytes (peri, colored red) and VAMCs (colored green). Dots represent expression levels in individual cells. (E, F) *Pdgfra*-positive cell transcriptomes were subsetted from the surface-associated and bone-associated datasets shown in (A, B) and merged together. Panel (E) shows the cluster UMAP, where transcriptionally distinct clusters are colored and numbered and (F) shows the dataset of origin for each transcriptome. CSA = cortical surface associated, BA = bone associated. **(G)** Gene ontology was performed on the differentially expressed genes identified in the comparison between leptomeningeal and dural cells as defined in (F) (> 1.3-fold average change, p adj < 0.05; see Extended Tables 1 and 2). Shown are selected categories (y axis) and the adjusted p-value (x-axis). The peach colored bars are categories enriched in leptomeningeal cells and the blue enriched in dural cells. **(H)** Single cell heatmap of select mRNAs that were differentially expressed (ExtendedTable 1) in the comparison between the leptomeningeal cells in clusters 0 and 2 and the dural cells in cluster 1 from the UMAPs in (E, F). Every row represents expression in an individual cell, and levels are colour-coded as per the adjacent key. **(I)** Violin plots showing relative expression levels of two mRNAs, *Ptgds* and *Igfbp7*, that were identified in the differential gene expression analysis as being highly-enriched in leptomeningeal (colored red) versus dural (colored blue) cells, respectively. **(J)** Transcriptional signatures for the putative leptomeningeal and dural cells were defined using the 50 most differentially-expressed mRNAs in the leptomeningeal versus dural cell comparison (Extended Table 1). Shown is a UMAP as in (E) highlighting cells expressing a transcriptional signature score of >0.5 for one or the other cell type.

In both preparations, this analysis (Fig. 1A, B; Fig. 1-1A-D) identified *Pdgfra*-positive mesenchymal cells, vasculature-associated cells including endothelial cells, pericytes and vasculature-associated mesenchymal cells (VAMCs), as well as various peripheral immune populations including macrophages, T cells and B cells. However, some cell types were specific to one or the other isolation. For the cortical surface analysis (Fig. 1A; Fig. 1-1A, B), we identified CNS cell types including astrocytes, microglia, a small number of neurons, and choroid plexus cells. By contrast, the bone-associated preparation (Fig. 1B; Fig. 1-1C, D) did not include CNS cell types or microglia, but did include Schwann cells deriving from cranial nerves (Bastedo et al., 2024; Carr et al., 2019).

We observed three types of *Pdgfrb*-positive mesenchymal cells within these datasets. In both cortical surface and bone-associated datasets there were *Pdgfrb*-positive pericytes and VAMCs that were negative for *Pdgfra* and positive for either the VAMC markers *Myh11* and *Acta2* or the pericyte markers *Rgs5* and *Kcnj8* (Fig. 1C, D; Fig. 1-1E, F). These vasculature mural cells are likely associated with meningeal blood vessels and parenchymal vasculature (Vanlandewijck et al., 2018). More importantly for our analysis, we also identified cells positive for *Pdgfrb*, *Pdgfra* and *Col1a1* that included putative dural meningeal cells in the case of the bone-associated dataset and leptomeningeal mesenchymal cells in the case of the cortical surface-associated dataset (Fig. 1C, D; Fig. 1-1E, F). Thus, our findings validated the dissections and identified likely meningeal mesenchymal cells (DeSisto et al., 2020; Pietilä et al., 2023; Vanlandewijck et al., 2018).

### Dural meninges and leptomeninges have distinct transcriptional signatures

We further analyzed the potential meningeal cells by subsetting, merging and reanalyzing *Pdgfra*-positive cells from both datasets. We performed one iteration of Harmony batch correction to ensure integration without over-correction (Korsunsky et al., 2019). UMAP-based cluster visualization (Fig. 1E) and analysis of the dataset of origin (Fig. 1F) identified two major groups of *Pdgra*-positive mesenchymal cells. Cluster 1 included cells almost exclusively from the bone-associated dataset, while clusters 0 and 2 included cells predominantly from the cortical surface dataset. We interpret these as bone-associated dural cells and cortical surface-associated leptomeningeal cells, respectively (DeSisto et al., 2020; Pietilä et al., 2023).

We directly compared these two populations using several approaches. First, we performed Pearson correlation analysis of averaged gene expression. The two mesenchymal cell populations were quite distinct with r = 0.844 (Fig. 1-1G). Second, we performed differential gene expression analysis (Extended Table 1) and identified 1239 differentially-expressed genes (average log2 fold change ≥ 1.3, adj p value < 0.5, expression in at least 10% of cells). 872 genes were enriched in the putative dural mesenchymal cells and 367 in the putative leptomeningeal cells. GO analysis (Kolberg et al., 2023) (Fig. 1G; Extended Table 2) showed that the dural mesenchymal cells were enriched for terms like extracellular matrix organization, skeletal system development, and ossification, consistent with their role as skull stromal cells. By contrast, the leptomeningeal cells were enriched for terms like active transmembrane transporter activity, cell junction organization and cell-cell signaling, consistent with their role as brain interface cells (Kolberg et al., 2023). Heatmaps and violin plots (Fig. 1H, I; Fig. 1-1H) confirmed enrichment for these mRNAs. The leptomeningeal cells were highly enriched for expression of the transmembrane transporter *Slc22a6*, the basal lamina component *Lama1*, the connexins *Gjb2* and *Gjb6*, and the prostaglandin D2 synthase *Ptgds*, which is associated with cerebral spinal fluid leak (Kondabolu et al., 2011). Conversely, the dural mesenchymal cells were enriched for *Igfbp7, Alpl, Chad, Ostn* and *Mgp,* all of which are implicated in osteochondrogenesis (Liu et al., 2023; Paracuellos et al., 2017; Sato et al., 2021; Zhang et al., 2018). We used these differentially-expressed genes to define a transcriptional signature that readily distinguished cluster 0/2 leptomeningeal cells versus cluster 1 dural mesenchymal cells (Extended Table 1). The leptomeningeal cell signature (Fig. 1J, blue) was comprised of 50 genes including *Ptgds, Slc22a6, Gjb6, Gjb2, Bmp6* and *Lama1*, while the dural mesenchymal cell signature (Fig. 1J, red) included 50 genes such as *Matn4, Igfbp7, H19, Clec11a* and *Smoc2*. Thus, both dura and leptomeninges are comprised of *Pdgfra*-positive mesenchymal cells, but the dural cells are more analogous to bone stromal cells while the leptomeningeal cells have a transcriptional profile consistent with a brain interface role.

### Brain interface astrocytes are transcriptionally distinct from cortical grey matter astrocytes

We next asked about cortical interface-associated astrocytes. Since the cortical surface scRNA-seq dataset contained relatively few astrocytes, we merged it with several additional astrocyte scRNA-seq datasets. First, we extracted cortical astrocyte transcriptomes from two recently-published scRNA-seq datasets, one including total postnatal day 60 (P60) adult cortex cells and the other P60 cortical grey matter cells (GEO GSM8072112; GSM8072111; Dennis et al., 2024) (Fig. 2-1A, B). Second, we took advantage of the recent finding that astrocytes at the brain surface (termed border astrocytes) are enriched for *Myoc* and *Gfap* mRNAs (Hasel et al., 2023). We therefore subsetted and included *Myoc*-expressing, *Gfap*-enriched astrocytes from a previously-published total brain astrocyte scRNA-seq dataset (Hasel et al., 2021; GEO GSE148611) (Fig. 2-1C, D). We merged all of these astrocyte transcriptomes with the astrocyte transcriptomes we had obtained in the cortical surface analysis (the dataset in Fig. 1A), and performed batch correction using Harmony.

**Figure 2.**
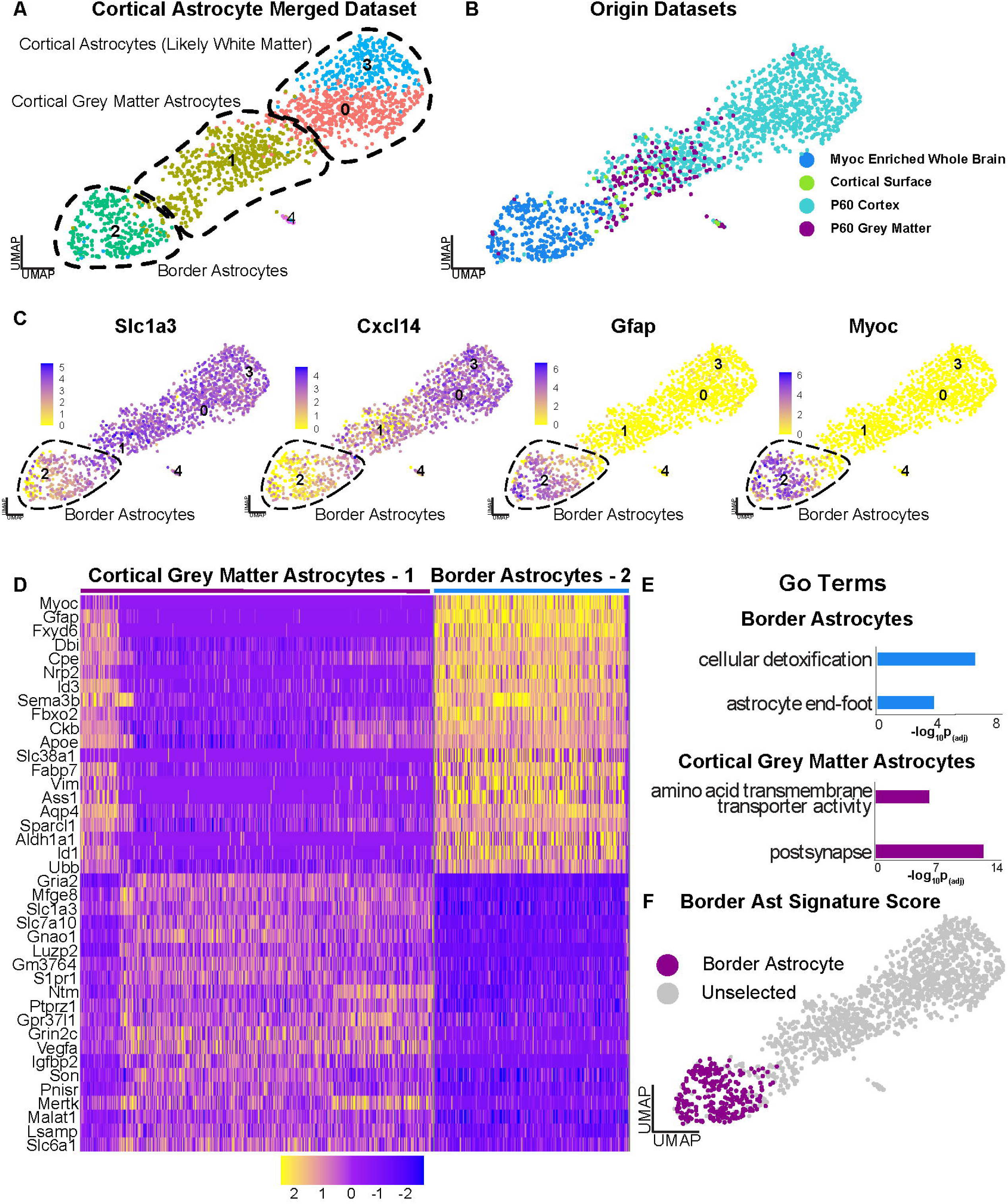
*Identification of transcriptionally-distinct border astrocytes by scRNA-seq analysis*. Also see Extended Figure 2-1 and Extended Tables 3 and 4. **(A, B)** Astrocyte transcriptomes were subsetted from 4 different murine scRNA-seq datasets; cortical surface-associated cells as in Figure 1A, P60 whole cortex and P60 cortical grey matter from Dennis et al., 2024 (GEO GSE255405), and whole brain astrocytes from Hasel et al., 2021 (GEO GSE148611). In the latter case, only astrocytes enriched for *Myoc* and *Gfap* were extracted (see Fig. 2-1C, D). These astrocyte transcriptomes were merged and batch-corrected using 1 iteration of Harmony. The resultant merged dataset is shown as UMAP visualizations with the transcriptionally-distinct clusters numbered and colour-coded in (A) and datasets of origin shown in (B). **(C)** UMAPs as in (A, B) overlaid for expression of two pan-astrocyte mRNAs, *Slc1a3* and *Cxcl14*, and two brain interface-enriched mRNAs, *Myoc* and *Gfap*. The putative border astrocytes are circled. Expression levels are color-coded as per the adjacent color keys. **(D)** Single cell heatmap of select mRNAs that were differentially expressed (Extended Table 3) in the comparison between the putative brain interface/border astrocytes (cluster 2 in panel A) and cortical grey matter astrocytes (cluster 1 in panel A). Every row represents expression in an individual cell, and levels are colour-coded as per the adjacent color key. **(E)** Gene ontology was performed on the differentially expressed genes identified in the comparison between border astrocytes and cortical grey matter astrocytes (> 1.3-fold average change, p adj < 0.05; see Extended Tables 3 and 4). Shown are selected categories (y axis) and the adjusted p-value (x-axis). **(F)** A transcriptional signature for the putative border astrocytes (cluster 2 in A) was defined using the 20 most differentially-expressed mRNAs in the border versus cortical grey matter astrocyte comparison (Extended Table 3). Shown is a UMAP as in (A) with cells expressing a border astrocyte transcriptional signature score >0.5 highlighted in purple.

This analysis (Fig. 2A, B) identified four astrocyte clusters in three distinct groups. One group of clusters (0 and 3) likely include cortical white matter astrocytes, since analysis of the dataset of origin showed they were almost completely comprised of astrocytes from the total cortex dataset, and did not include grey matter transcriptomes. The grey matter transcriptomes were instead in cluster 1, which also included transcriptomes from the total cortex, and thus likely includes cortical parenchymal astrocytes. The final cluster (2) was largely comprised of *Myoc*-enriched whole brain astrocytes, as well as cells from the cortical surface dataset, suggesting that they were border astrocytes. All cells expressed pan-astrocyte markers such as *Slc1a3* (GLAST) and *Cxcl14* and, as predicted, cluster 2 was enriched for *Myoc* and *Gfap* (Fig. 2C) (Batiuk et al., 2020).

To better-understand the putative cluster 2 border astrocytes, we directly compared them to cluster 1 cortical parenchymal/grey matter astrocytes using differential gene expression analysis. 575 mRNAs differed significantly between the two groups (average log2 fold change ≥ 1.3, adj p value < 0.5, expression in at least 10% of cells) (Extended Table 3). Cluster 2 border astrocytes were enriched for 381 mRNAs including *Myoc*, *Gfap, Fbxo2, Id1, Vim* and *Agt*, as illustrated in a heatmap (Fig. 2D; Extended Table 3). Gene ontology analysis of these differentially-enriched border astrocyte genes (Extended Table 4) identified terms that included cellular detoxification and astrocyte end-foot (Fig. 2E). By contrast, cluster 1 grey matter astrocytes were enriched for 194 mRNAs, including *Gria2, Igfbp2*, and *Slc1a3* (Fig. 2D; Extended Table 3) and gene ontology identified terms such as post-synapse and amino acid transmembrane transporter (Fig. 2E; Extended Table 4). We used these differentially-expressed genes to define a specific transcriptional signature for the putative *Myoc*-positive border astrocytes comprised of 20 genes including *Myoc, Gfap, Fbxo2, Slc38a1* and *Fxyd6* (Fig. 2F; Extended Table 3).

### Visualization of leptomeningeal:astrocyte interactions at the cortical brain interface

One caveat of the scRNA-seq data is that it does not provide spatial information about the interface cells. We therefore performed two complementary morphological studies, lineage tracing plus immunostaining and single cell spatial transcriptomics. For the lineage tracing analysis, we initially used a well-characterized mouse line where CreERT2 is driven from the endogenous *Pdgfra* locus (Kang et al., 2010). We crossed these to another mouse line carrying a floxed TdTomato transgene in the *Rosa26* locus; when these crossed *Pdgfra-CreERT2;R26RtdT* mice are exposed to tamoxifen, TdTomato will be expressed in and specifically tag *Pdgfra*-positive mesenchymal cells. *Pdgfra*-positive OPCs will also be genetically-tagged, but these are readily distinguished by their location, morphology and transcriptional profiles. We therefore exposed crossed mice to tamoxifen for 5 days, 10 days later isolated the cortex and overlying skull, decalcified the skull, and analyzed sagittal sections from layer 1 through to the top of the skull (Fig. 3A). This analysis (Fig. 3B) showed that as predicted both the dura and the leptomeninges were TdTomato-positive and that the leptomeninges were only two to three layers thick in this region. Moreover, immunostaining for GFAP (Fig. 3B), which is enriched in the border astrocytes (Fig. 2C), showed that the TdTomato-positive leptomeningeal cells were immediately adjacent to GFAP-positive astrocytes. Similar findings were made by immunostaining sections for PDGFRα protein and GFAP (Fig. 3-1A).

**Figure 3.**
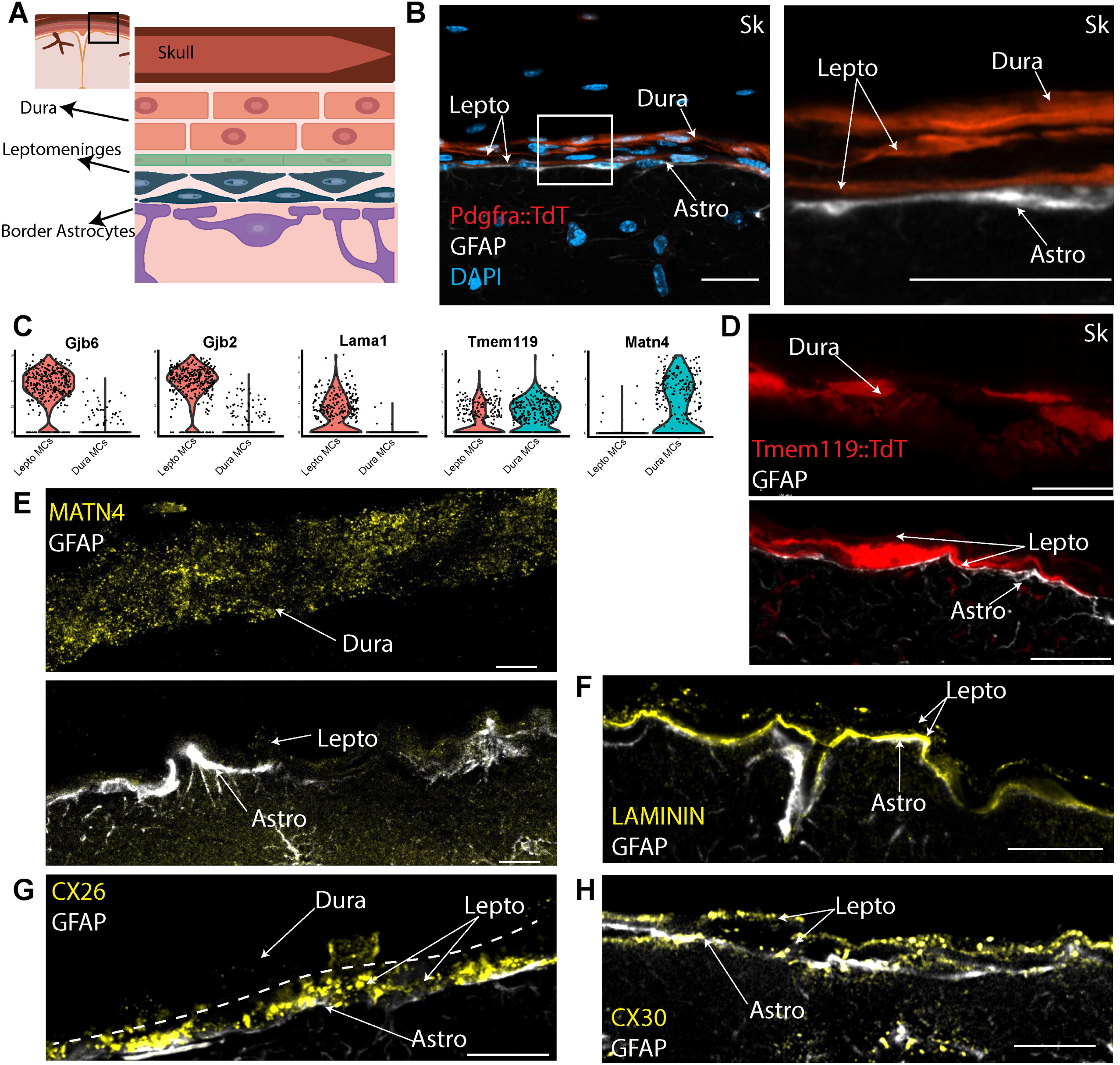
*Lineage tracing and immunostaining identify the leptomeninges and the dura at the cortical brain interface*. Also see Extended Figure 3-1. **(A)** Schematic showing the region of the cortex and overlying skull that were analyzed in the lineage tracing and immunostaining studies. **(B)** Ten week old *Pdgfra-CreERT2;R26RtdT* mice were treated with tamoxifen and tissues collected 10 days after the final administration. The brain and skull were decalcified and sagittal cortical sections were immunostained for GFAP (white) and counterstained with DAPI (blue). Shown is a representative confocal image of the brain interface and skull showing TdTomato (red), GFAP (white), and DAPI (blue). The boxed region is shown at higher magnification to the right. Lepto = leptomeninges, Astro = border astrocytes, Sk = skull. **(C)** Violin plots showing the relative expression of *Gjb6, Gjb2, Lama1, Tmem119,* and *Matn4* in the leptomeningeal (colored red) versus dural (colored blue) cells shown in Figure 1E, F. Dots represent individual transcriptomes. **(D)** Ten week old *Tmem119CreErt2;R26RtdT* mice were treated with tamoxifen and tissues collected 10 days later. The brain and skull were decalcified and sagittal cortical sections immunostained for GFAP (white). Shown is a representative confocal image of the cortical interface showing TdTomato (red) and GFAP (white). Both upper and lower panels come from the same field of view, but are shown separately since the dura/skull was separated from the brain interface. Lepto = leptomeninges, Astro = border astrocytes, Sk = skull. **(E)** High magnification confocal images of a sagittal section through the brain interface and skull, immunostained for GFAP (white) and MATN4 (yellow). Both upper and lower panels come from the same field of view, but are shown separately since the dura/skull was separated from the brain interface. Lepto = leptomeninges, Astro = border astrocytes. **(F)** High magnification confocal image of a sagittal adult cortical interface section immunostained for GFAP (white) and Laminin (yellow). Note that the Laminin immunoreactivity is localized between the border astrocytes and the leptomeningeal cells. Lepto = leptomeninges, Astro = border astrocytes. **(G)** High magnification confocal image of a sagittal adult cortical interface immunostained for GFAP (white), and Cx26 (yellow). The hatched line delineates the border between the leptomeninges and the dura. Lepto = leptomeninges, Astro = border astrocytes. **(H)** High magnification confocal image of a sagittal adult cortical interface immunostained for GFAP (white), and Cx30 (yellow). Lepto = leptomeninges, Astro = border astrocytes. For all panels, scale bars = 10um.

We performed a similar analysis with a *Tmem119-CreERT2;R26RtdT* mouse line that has been widely used to label microglia (Kaiser and Feng, 2019), based upon our finding that *Tmem119* is expressed in both the dura and leptomeninges (Fig. 3C). As predicted, following tamoxifen treatment, both the dura and leptomeninges were TdTomato-positive in the *Tmem119-CreERT2;R26RtdT* mice (Fig. 3D), as were microglia located in the cortical parenchyma. As seen with the *Pdgfra-CreERT2*-based lineage tracing, the leptomeninges were only several cell layers thick at this point above the cortex. Moreover, immunostaining for GFAP showed that the GFAP-positive border astrocytes were closely juxtaposed to the TdTomato-positive leptomeningeal cells (Fig. 3D).

Both the dura and leptomeninges express *Pdgfra* and *Tmem119*, and thus these lineage tracing approaches do not distinguish the two mesenchymal compartments (Pietilä et al., 2023). We therefore immunostained for proteins predicted to be differentially-expressed in the transcriptional data. For the dura, we immunostained decalcified sagittal cortical sections that included the skull for MATN4, which is highly enriched in the putative dural mesenchymal cells (Fig. 3C). For the leptomeninges, we immunostained similar sections for Laminin (*Lama1*), or the gap junction proteins Connexin 26 (*Gjb2*) or Connexin 30 (*Gjb6*), all of which are highly-enriched in the leptomeningeal cells (Fig. 3C) (DeSisto et al., 2020; Pietilä et al., 2023). Of these, *Gjb2* is also expressed at low levels in border astrocytes and *Gjb6* is widespread in astrocytes (Fig. 3-1B-D). In all cases we double-labelled sections for GFAP to visualize the border astrocytes. As predicted, immunoreactivity for the extracellular matrix protein MATN4 was not detectable in the leptomeninges but was observed in a punctate pattern in the dura (Fig. 3E). By contrast, Laminin, Cx26/Gjb2 and Cx30/Gjb6 were detectable in the leptomeninges but not the dura (Fig. 3F-H). Laminin immunoreactivity was specifically associated with the basal lamina between the GFAP-positive border astrocytes and the leptomeningeal cells, while Cx26 and Cx30 immunoreactivity were present in a punctate pattern in the leptomeningeal cells immediately adjacent to GFAP-positive border astrocytes. As predicted (Fig. 3-1B, D) Cx30 mRNA was also detected in border astrocytes, as was Cx26 at apparently lower levels (Huang et al., 2021; Lynn et al., 2011; Nagy et al., 2011). These studies thus identify new lineage tracers for the leptomeninges, validate our findings from the scRNA-seq datasets and highlight the close interactions between border astrocytes and leptomeningeal cells at the cortical brain interface.

### Single cell spatial transcriptomics to characterize the cortical surface interface

The lineage tracing and immunostaining studies are limited by their ability to only analyze several marker genes/proteins at a time. To obtain a more global overview of the cortical brain interface we performed single cell multiplexed *in situ* gene expression analysis using the Xenium platform (Janesick et al., 2023; Willis et al., 2025). We did this with two different probesets, one a custom 480 gene probeset optimized to distinguish different mesenchymal cell populations, including those of the meninges, and the second a standard Xenium brain probeset with a custom 100 gene add-on comprised of probes allowing better resolution of non-neural brain cell types (see Materials and Methods; Extended Table 9).

We used these probesets to analyze the leptomeninges and the adjacent cortical layer one in coronal forebrain sections from freshly-frozen adult murine brain tissue (see Fig. 4A). For sections analyzed using the mesenchymal probeset, we also included the dura. We then performed Xenium-based multiplexed *in situ* gene expression analysis as described (Dennis et al., 2024; Willis et al., 2025) using one of the two probesets. For the mesenchymal probeset, we analyzed 6 sections in total from 4 different mice. For the brain probeset, we ran 1 section each from 2 different adult mouse brains. We augmented this with a recently-published Xenium dataset obtained using the same brain probeset to analyze 3 coronal sections of the adult cortex at a similar level from 3 different mice (Willis et al. 2025; GEO GSE266689). In all cases we defined a region of interest (ROI) that spanned the midline and included cortical layer 1, the top of adjacent layer 2 plus the overlying leptomeninges (see Fig. 4A). After initial processing of data from individual sections, we excluded cells where the detected genes and total transcript counts per cell were +/-2.5 or more standard deviations from the mean in either parameter since these were likely cellular fragments or cellular doublets. We then merged cells from different sections/animals analyzed using the same probeset.

**Figure 4.**
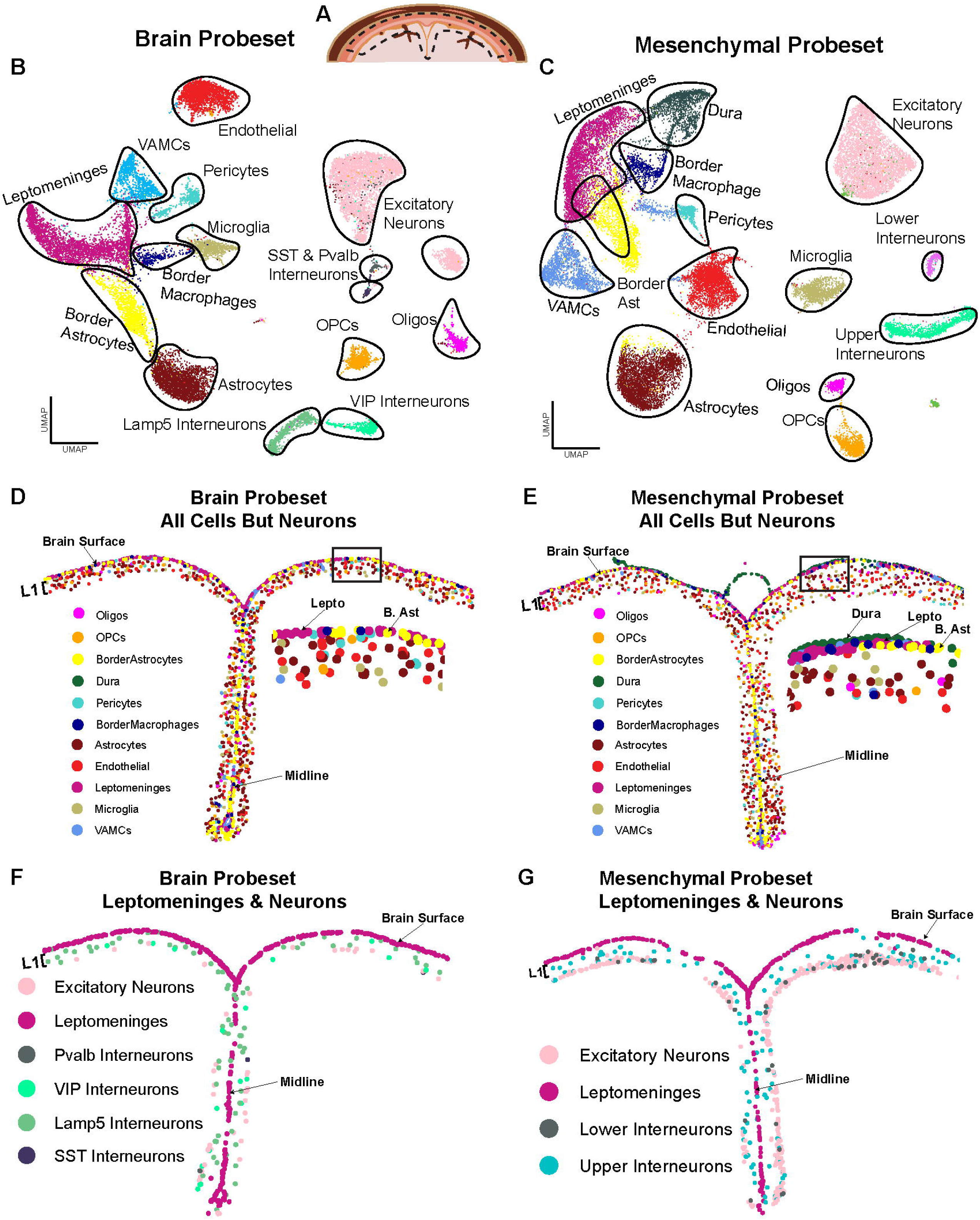
*Single cell multiplexed in situ gene expression analysis with two different probesets identifies the spatial location of cortical interface and layer one cell types*. Also see Extended Figure 4-1 and Extended Table 9. Coronal sections of adult mice were analyzed by Xenium-based single cell multiplexed *in situ* gene expression analysis with either a brain-targeted panel probing 347 genes or a mesenchymal cell-targeted panel probing 480 genes (see Extended Table 9). The ROI that was analyzed is shown in (A). Datasets from different sections and conditions were then merged and cell types identified by marker gene expression. **(A)** Schematic of a coronal section through the adult cortex in the midline region showing the ROI that was analyzed (outlined with hatched black lines). **(B, C)** UMAP cluster visualization of the merged transcriptomes from analysis with the brain-targeted probeset (B) or the mesenchymal cell-targeted probeset (C), annotated for cell types. Each dot represents a single cell. VAMC = vasculature-associated mesenchymal cell, Oligos = oligodendrocytes, SST = somatostatin, Pvalb = parvalbumin, VIP = vasoactive intestinal peptide. **(D, E)** Spatial plots of the midline and adjacent cortical interface and layer one from the ROI of representative sections analyzed using the brain (D) or mesenchymal (E) probesets, showing all cell types except neurons, color-coded as per the legend. The boxed regions are also shown in enlarged views. Lepto = Leptomeninges, B. Ast = Border astrocytes. **(F, G)** Spatial plots as in (D, E) from the same representative sections analyzed using the adult brain (F) or adult mesenchymal (G) probesets showing the leptomeninges and all of the neurons that were identified, color-coded. L1 denotes cortical layer one.

With the mesenchymal/non-neural cell probeset, we ultimately obtained 29644 total cellular transcriptomes from the merged ROIs of 6 sections/4 brains and with the brain probeset, we obtained 20371 transcriptomes from merged ROIs of 1 section each from 5 brains. With both probesets, transcriptomes from the different sections/brains were well-integrated, as shown on UMAPs (Fig. 4B, C; Fig. 4-1A, B). With two exceptions, both probesets defined the same cell types, as determined by analysis of a panel of marker genes (see Materials and Methods) (Fig. 4B, C; Fig. 4-1C, D). The exceptions were dural cells, which as predicted were only present in the mesenchymal panel sections, and interneurons (discussed further below).

Spatial plots identified the locations of the different identified cell types (Fig. 4D-G). The brain interface was comprised of leptomeningeal cells, macrophages and border astrocytes that were tightly packed, interspersed with vasculature-associated cells (mural cells and endothelial cells) presumably deriving from penetrating blood vessels. Layer one was instead relatively cell-sparse, and included astrocytes, oligodendrocytes, oligodendrocyte precursor cells (OPCs) and vasculature-associated cells. As predicted, the only neurons in layer one were interneurons (Fig. 4B, F). The brain panel defined these as expressing *Lamp5* or *Vip* (Fig. 4B, F), consistent with previous immunocytochemical analyses (Gesuita and Karayannis, 2021; Schuman et al., 2021, 2019; Yao et al., 2021). The mesenchymal panel instead defined two interneuron clusters that differed in their spatial locations (from hereon called upper or lower interneurons) (Fig. 4C, G). The border of layer one was clearly delineated by layer 2 excitatory neurons and a few *Somatostatin* or *Parvalbumin*-expressing interneurons that were included in the ROI (Fig. 4F, G), again as previously reported (Lim et al., 2018; Tremblay et al., 2016; Yao et al., 2021). We used these datasets to analyze the brain interface and layer one cellular neighborhoods in greater detail. Consistent with the immunostaining analyses (Fig. 3), the leptomeninges at the brain interface were a continuous layer of cells that were lined by border astrocytes on the CNS side and by the dura on the skull side (Fig. 5A). Resident macrophages were scattered along this interface. These are likely analogous to the previously-described border macrophages (Jordão et al., 2019; Kolabas et al., 2023; Pietilä et al., 2023).

**Figure 5.**
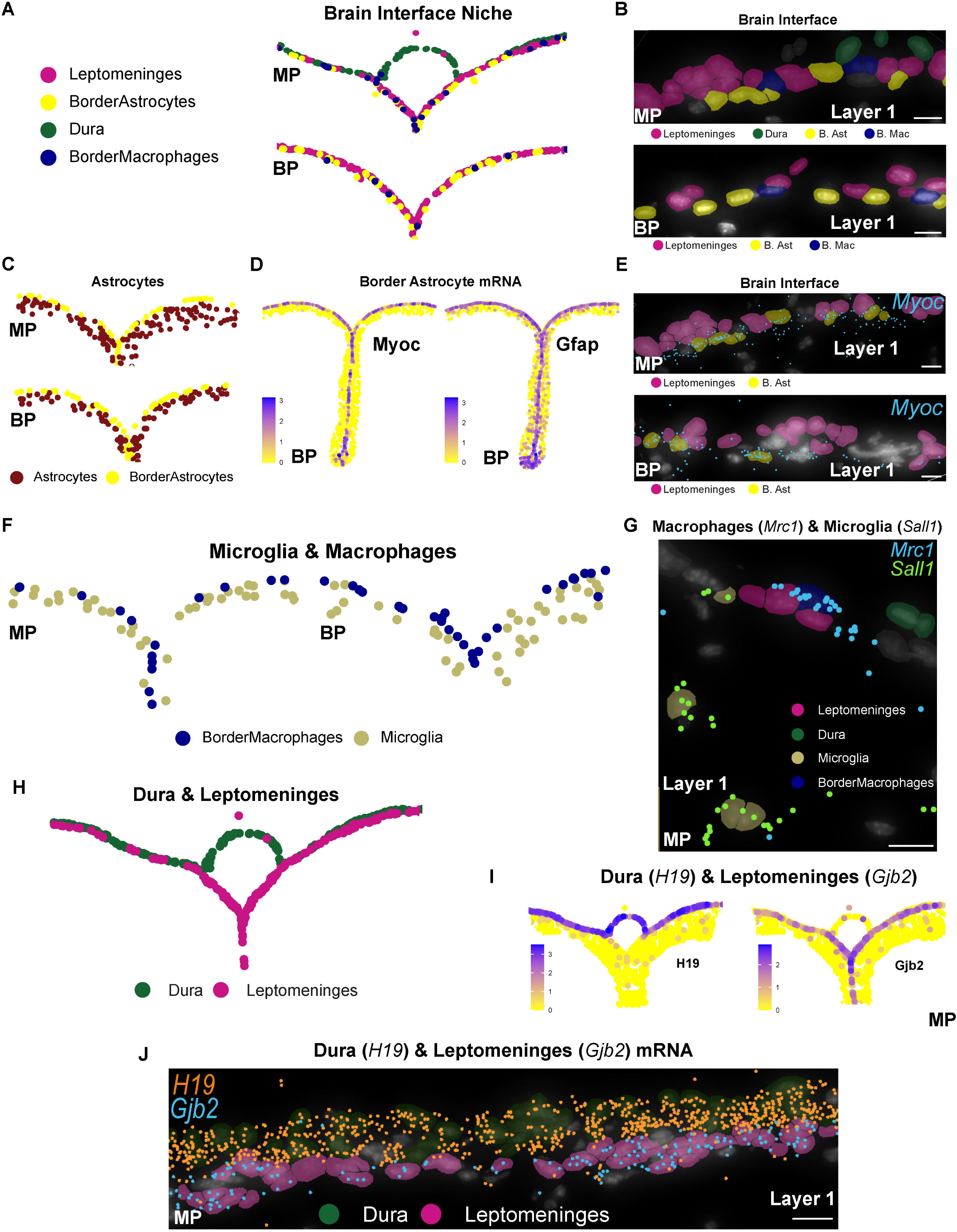
*Single cell spatial transcriptomic analysis defines the cortical brain interface structure and cell types*. Also see Extended Table 5. Coronal sections of adult mice were analyzed by Xenium-based single cell multiplexed *in situ* gene expression analysis with either a brain-targeted probeset for 347 genes or a mesenchymal cell-targeted probeset for 480 genes (see Extended Table 9). The ROI and UMAPs of the resultant merged datasets are shown in Fig. 4A-C. **(A)** Spatial plots of the cortical interface analyzed with the mesenchymal (MP, top) or brain (BP, bottom) probesets showing leptomeningeal cells (pink), border astrocytes (yellow), border macrophages (dark blue) and dural cells (green, mesenchymal probeset only). **(B)** High resolution Xenium Explorer images of the cortical interface region analyzed with the mesenchymal (MP, top) or brain (BP, bottom) probesets showing the spatial arrangement of leptomeningeal cells (pink), border astrocytes (yellow), border macrophages (blue) and dural cells (green, mesenchymal probeset only). **(C)** Spatial plots of the cortical interface and layer one analyzed with the mesenchymal (MP, top) or brain (BP, bottom) probesets showing border astrocytes (yellow) and parenchymal astrocytes (brown). **(D)** Spatial plots of *Myoc* and *Gfap* mRNA expression in all cells of the full ROI analyzed using the brain probeset (BP). Relative expression levels are coded as per the adjacent keys. **(E)** High resolution Xenium Explorer image of the cortical interface analyzed with the mesenchymal (MP, top) or brain (BP, bottom) probesets showing expression of *Myoc* mRNA (light blue dots) relative to the leptomeningeal cells (pink) and border astrocytes (yellow). Nuclei of other cell types (white) are also shown. **(F)** Spatial plots of the cortical interface and layer one analyzed with the mesenchymal (MP, left) or brain (BP, right) probesets showing border macrophages (blue) and microglia (tan). **(G)** High resolution Xenium Explorer image of the cortical interface and layer one analyzed with the mesenchymal probeset (MP) showing expression of *Mrc1* mRNA (light blue dots) and *Sall1* mRNA (bright green dots) relative to the border macrophages (dark blue), microglia (olive) and leptomeningeal cells (pink). Nuclei of other cell types (white) are also shown. **(H)** Spatial plot of the cortical interface and midline analyzed with the mesenchymal probeset showing dural (green) and leptomeningeal (pink) cells. **(I)** Spatial plots of the dural mRNA *H19* (left) or the leptomeningeal mRNA *Gjb2* (right) in all cells at the cortical interface, midline and layer one analyzed using the mesenchymal probeset (MP). Relative mRNA expression levels are coded as per the adjacent keys. **(J)** High resolution Xenium Explorer image of the cortical interface region analyzed with the mesenchymal probeset (MP) showing expression of *H19* mRNA (orange dots) and *Gjb2* mRNA (bright blue dots) relative to the leptomeningeal cells (pink) and dural cells (green). Nuclei of other cell types (white) are also shown. Low magnification spatial plots in A, C, D, F, H, and I were generated using Seurat and show part of the ROI centered on the cortical midline region from one representative section each, with each dot representing the centroid of one cell. High resolution spatial plots (B, E, G and J) were generated using Xenium Explorer. Scale bars = 10um.

A higher resolution analysis highlighted the close association between these different cell types (Fig. 5B). The cortical surface leptomeninges were 2 to 3 cells wide, with the lowest layer immediately adjacent to the border astrocytes. The border macrophages were interspersed within this leptomeningeal layer. Each of these interface-associated cell types differed transcriptionally from similar cells within other regions of the meningeal/cortical layer one tissue. Specifically, the border astrocytes were transcriptionally-distinct and spatially segregated from their parenchymal astrocyte counterparts in layer one (Fig. 5C), and, consistent with the scRNA-seq, were enriched for *Myoc* and *Gfap* (Fig. 5D, E). The border macrophages were also transcriptionally and spatially-distinct from the parenchymal microglia (Fig. 5F). The border macrophages were closely-associated with the leptomeningeal cells, with the occasional macrophage in the parenchyma where they were associated with vasculature cells and were presumably present in blood vessels.

To confirm these macrophage/microglia cell type assignments, we performed differential gene expression (Extended Table 5) on the macrophages and microglia from the cortical surface-associated scRNA-seq data (Fig. 1A). This analysis showed that surface macrophages versus microglia were enriched for *Mrc1* versus *Sall1*, respectively, as previously reported (Jurga et al., 2020; Paolicelli et al., 2022). Since probes for both of these genes were present in the mesenchymal probeset, we analyzed their expression spatially. This analysis confirmed that the cells identified as border macrophages were positive for *Mrc1* while those identified as microglia were specifically positive for *Sall1* (Fig. 5G).

These analyses also supported several conclusions regarding the meningeal mesenchymal cells. First, the dural mesenchymal cells were readily distinguished from the leptomeningeal cells; they were located on the skull side of the leptomeninges (Fig. 5H) and expressed mRNAs such as *Matn4* and *H19* but not the leptomeningeal mRNA *Gjb2* (Fig. 5I, J). Second, the leptomeningeal cells segregated into two distinct clusters on higher resolution UMAPs of both the brain and mesenchymal Xenium datasets (Fig. 6-1A, B). One of these two clusters was differentially enriched for *Lama1* and *Slc22a6* mRNAs (Fig. 6A). We have called these cells lower leptomeningeal cells since on spatial plots they extend down the midline and are immediately adjacent to the cortex (Fig. 6B-D), and since Laminin protein is specifically enriched adjacent to the border astrocytes (Fig. 3F). The second cluster was instead enriched for *Dpp4* and *Cdh1* (Fig. 6A). We have called these cells upper leptomeningeal cells since they do not extend down the midline and are superficial to the lower leptomeningeal cells (Fig. 6B-D). It is likely that lower leptomeningeal cells are pial cells and the upper arachnoid/arachnoid barrier cells since (i) pial but not arachnoid cells extend down the midline, (ii) pial cells express *Lama1* to form the basal lamina, and (iii) arachnoid barrier cells have been reported to express *Dpp4* (DeSisto et al., 2020; Pietilä et al., 2023).

**Figure 6.**
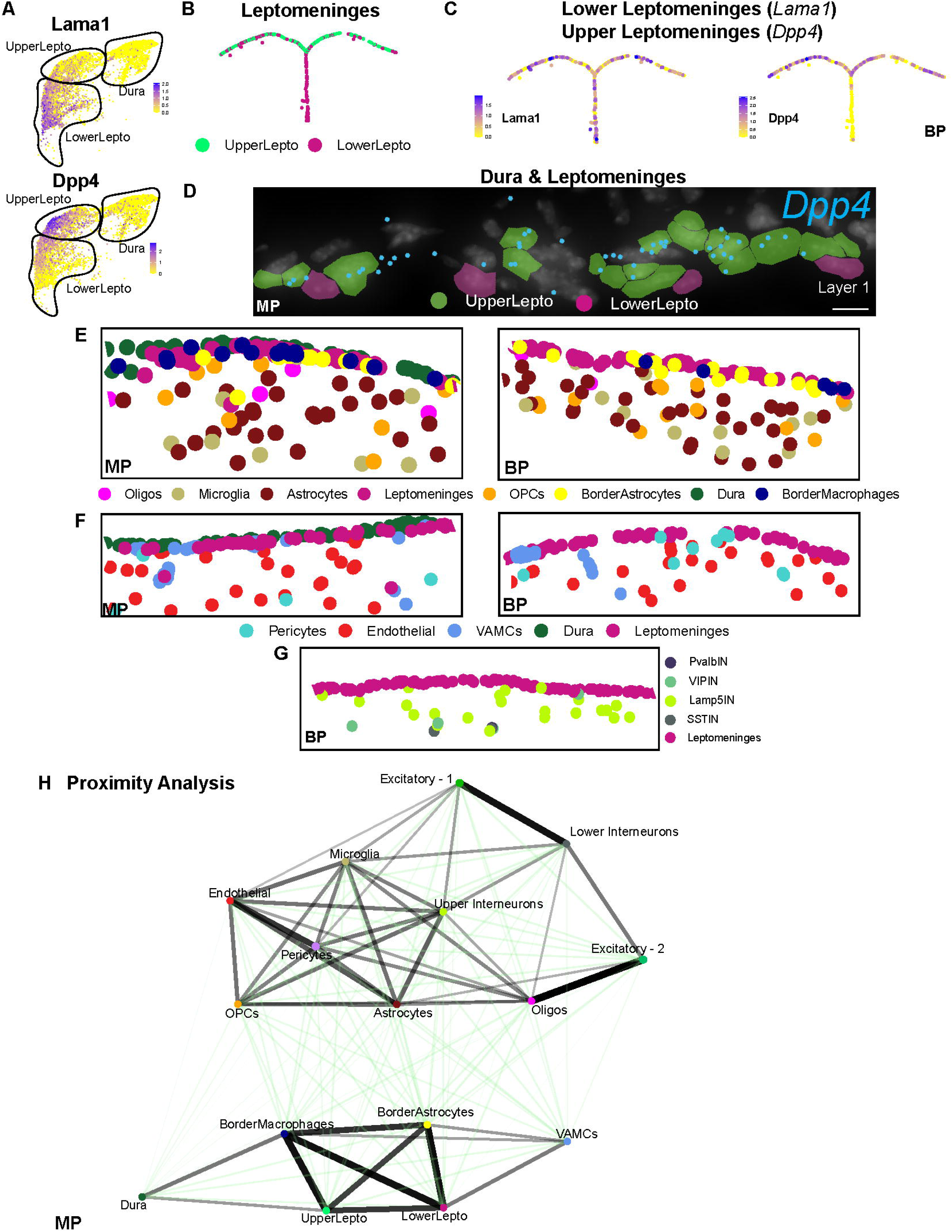
Single cell spatial transcriptomic analysis defines proximal cellular neighborhoods for the cortical brain interface and layer one. Also see Extended Figure 6-1. Coronal adult cortical sections were analyzed by Xenium-based single cell multiplexed *in situ* gene expression analysis with either a brain-targeted probeset for 347 genes or a mesenchymal cell-targeted probeset for 480 genes (see Extended Table 9). The ROI and UMAPs of the resultant merged datasets are shown in Fig. 4A-C. **(A)** UMAP cluster visualization of the *Pdgfra*-positive mesenchymal cell clusters from the mesenchymal probeset dataset shown in Fig. 4C, overlaid for *Lama1* and *Dpp4* mRNAs, which distinguish the upper and lower leptomeningeal clusters. The lines outline the three clusters defined at higher resolution (Extended Figure 6-1A) and are annotated for mesenchymal cell type. Gene expression levels are color-coded as per the adjacent keys. **(B)** Spatial plot of the cortical interface and midline analyzed with the brain probeset (BP) showing upper (bright green) and lower (pink) leptomeningeal cells. Note the lower but not upper leptomeningeal cells extend down the midline. **(C)** Spatial plots of the lower leptomeninges mRNA *Lama1* (left) or the upper leptomeningeal mRNA *Dpp4* (right) in total leptomeningeal cells at the cortical interface and midline analyzed using the brain probeset (BP). Relative mRNA expression levels are coded as per the adjacent keys. **(D)** High resolution Xenium Explorer image of the cortical interface region analyzed with the mesenchymal probeset (MP) showing expression of *Dpp4* mRNA (blue dots) relative to the upper leptomeningeal (green) and lower leptomeningeal (pink) cells. Nuclei of other cell types (white) are also shown. Scale bar = 10um. **(E)** Spatial plots of the cortical interface and layer one analyzed with the mesenchymal (MP, left) and brain (BP, right) probesets showing the oligodendrocytes (pink), microglia (tan), parenchymal astrocytes (dark brown), leptomeninges (dark pink), OPCs (orange), border astrocytes (yellow), dural cells (green, mesenchymal probeset only), and border macrophages (dark blue). **(F)** Spatial plots of the cortical interface and layer one analyzed with the mesenchymal (MP, left) and brain (BP, right) probesets showing the pericytes (turquoise), endothelial cells (red), VAMCs (blue), dural cells (green, mesenchymal probeset only), and leptomeninges (dark pink). **(G)** Spatial plot of the cortical interface and layer one analyzed with the brain probeset (BP) showing the leptomeninges (dark pink) and interneurons expressing *Lamp5* (lime green, Lamp5IN), *Vip* (blue-green, VIPIN), *Somatostatin* (grey, SSTIN) or *Parvalbumin* (black, PvalbIN). **(H)** Proximity analysis showing the relative strength of statistically-significant interactions between different cell types located in the cortical brain interface and adjacent layer one, as analyzed from the dataset obtained using the mesenchymal probeset. The nodes are color-coded and annotated for cell type, and the weight of the line indicates the strength of the interaction. Black lines indicate significant interactions based on a permutation test, comparing our data to a null distribution created from the random permutation of cell labels with fixed positions (See Methods and Materials for details). Green lines indicate interactions that were not statistically significant. Excitatory neuron-1 and excitatory neuron-2 denote transcriptionally-distinct excitatory neuron clusters. Cell centroid information was used to perform this analysis, with a proximal interaction being defined as one where cell centroids were within 70µm of another cell centroid. This unsupervised analysis identifies two neighborhoods, the cortical brain interface and layer one, as well as a partial neighborhood centered around the layer 2 excitatory neurons at the boundary of layer one. Oligos = oligodendrocytes, Lepto = leptomeningeal cells. Low magnification spatial plots (B, C, E-G) were generated using Seurat and each dot represents the centroid of one cell. The high resolution spatial plot in (D) was generated using Xenium Explorer and the proximity map in (H) was generated using Giotto.

### Spatial characterization of the layer 1 cellular environment

We also used these datasets to more precisely define the layer 1 cellular environment. Spatial plots showed that layer one was relatively cell-sparse, and included scattered microglia, astrocytes, OPCs and oligodendrocytes as well as vasculature-associated endothelial and mural cells (Fig. 6E, F). As indicated above, both probesets identified interneurons but not excitatory neurons in layer one, as compared to layer 2, which was densely packed with both excitatory and inhibitory neurons (Fig. 4C, D; data not shown). The brain probeset identified a more abundant *Lamp5*-expressing interneuron population localized throughout layer one, and a second less abundant *Vip*-expressing population largely located closer to the layer 2 border (Fig. 6G). As predicted, this analysis also identified interneurons in layer 2 but not layer 1 that expressed *Sst* or *Parvalbumin* (Fig. 6G) (Lim et al., 2018; Yao et al., 2021).

### Brain interface and layer one cells comprise different cellular neighborhoods

We next defined the cellular environment of the brain interface itself and the adjacent cortical layer 1, initially by defining cells that were in close proximity to each other using *cellProximityEnrichment* in Giotto v. 4.0.2 (described in detail in Dries et al. 2021) to analyze the single cell spatial transcriptomic data. We defined cells as being proximal to each other if the cell centroids were within 70 µm of each other, and termed each proximal association an interaction. We obtained similar results with both probesets (Fig. 6H; Fig. 6-1C), identifying two distinct neighborhoods. One of these included the brain interface cells; the upper and lower leptomeninges, border astrocytes and border macrophages, all of which were located in very close proximity to each other. Also in the vicinity were the dural cells and the vascular-associated mesenchymal cells that are commonly associated with larger blood vessels. By contrast, the brain interface cells were largely not significantly associated with other layer one cell types, although spatial plots did detect the occasional neural cell close to border astrocytes at the interface (for example, see Fig. 6E). Nonetheless, of total heterocellular interactions with border astrocytes, only 16.5% involved cells other than leptomeninges, border macrophages and vasculature cells.

The second proximal neighborhood included the cortical layer 1 and layer 2 cells other than border astrocytes; *Lamp1*-positive upper interneurons, OPCs, oligodendrocytes, microglia, parenchymal astrocytes, excitatory neurons at the layer one/two border, as well as the endothelial cells and pericytes associated with smaller vessels and capillaries. Except for associations involving layer 2 neurons, none of these interactions were statistically as robust as those between the border interface cells, consistent with the relatively sparse cellular density in layer 1 (Schuman et al., 2021). Thus, the cellular environments of the brain interface and cortical layer 1 are distinct, and the brain interface environment is likely predominantly determined by the border astrocytes, border macrophages, and leptomeningeal cells.

### Leptomeningeal cells express ligands predicted to act on both border astrocytes and macrophages

We defined ligands made within the brain interface that might influence the biology of this cell dense neighborhood. To do this, we used the scRNA-seq datasets (Fig. 1A, B) and a previously-published ligand-receptor database (Toma et al., 2020). We defined a ligand mRNA as expressed if it was detectable in at least 5% of the relevant cell type. We identified 65 leptomeningeal ligands, 48 border macrophage ligands and 54 border astrocyte ligands (Extended Table 6). Of these, 15 were expressed by all three cell types, including ligands such as *Bmp6, Efnb1, Hbegf, Il18, Vegfb* and *Gas6*. Fifteen were expressed by leptomeninges and border astrocytes, including *Ctf1* and *Gdf11*, 12 by leptomeninges and border macrophages, including *Wnt5a, Igf2* and *Tgfb1*, while one, *Sema4d*, was expressed by border astrocytes plus macrophages. The remaining ligands were expressed by only one of the three cell types; leptomeninges expressed 13 distinct ligands including *Bmp4, Bmp5, Bmp7, Sema3a* and *Igf2,* border macrophages 9 distinct ligands including *Osm, Tnf*, and *Il1b*, while border astrocytes specifically expressed 13 ligands including *Fgf1, Tgfb2, Wnt7a* and the semaphorins *Sema3b, Sema4g* and *Sema5b*.

To ask if these ligands might influence the brain interface neighborhood, we used the same scRNAseq data and ligand-receptor database to define receptors expressed by each interface cell type. This analysis defined 102, 95 and 79 receptors expressed by the leptomeninges, border astrocytes and border macrophages respectively (Extended Table 7). We used this information to predict which ligands might be biologically-active. This modeling (Fig. 7A, B; Extended Table 9) predicted 78 ligand-receptor interactions within the brain interface neighborhood. Most ligands (52) had receptors on all three brain interface cell types, but the remainder were more specific.

**Figure 7.**
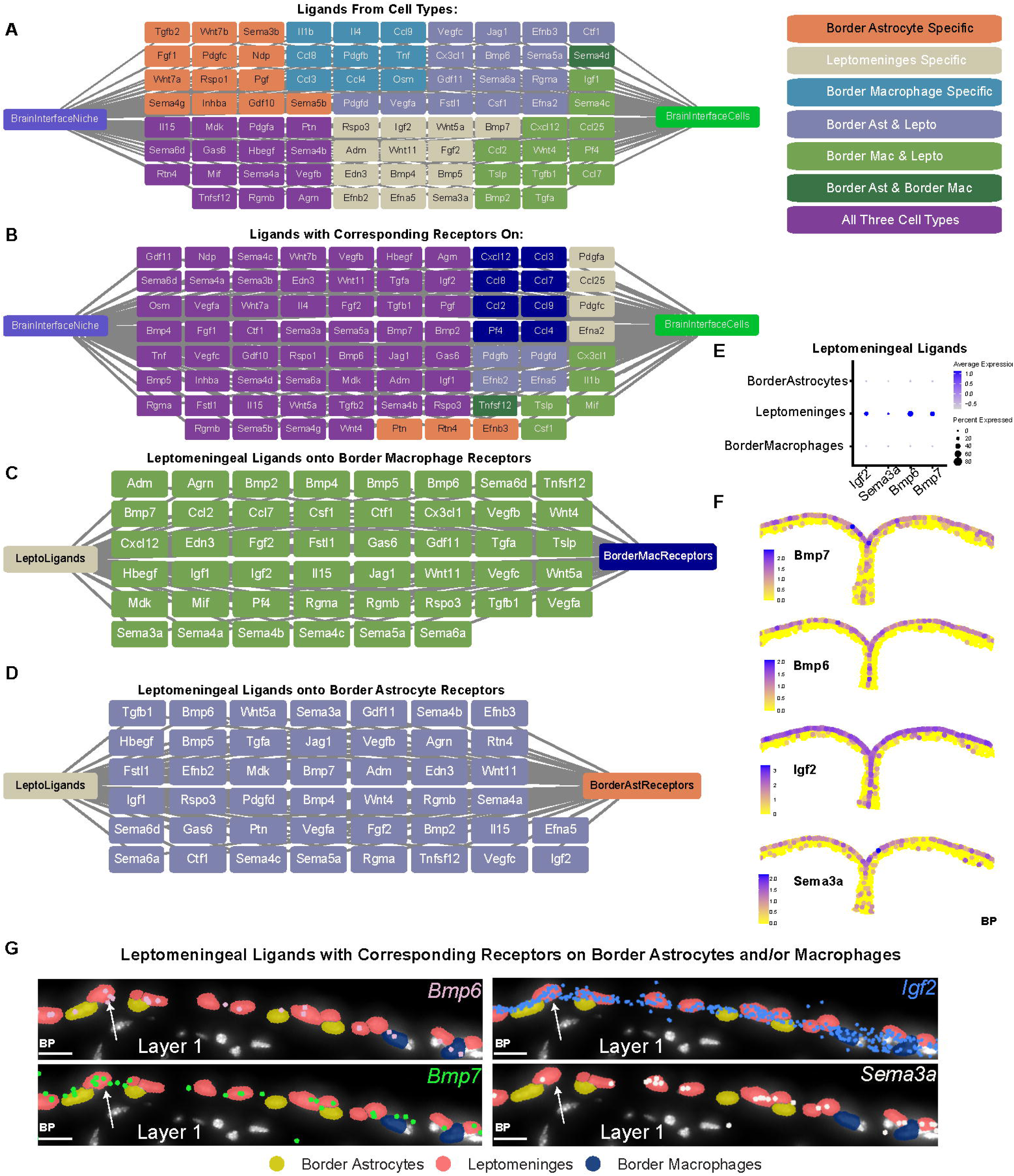
*Leptomeningeal cells express ligands predicted to regulate border astrocytes and border macrophages*. Also see Extended Figure 7-1 and Extended Tables 6-8. **(A-D)** Leptomeningeal cells, border astrocytes and border macrophages were analyzed for their expression of ligand and ligand receptor mRNAs using the scRNA-seq data (see Extended Tables 6-8). Ligand or receptor mRNAs were included if they were expressed in ≥ 5% of the relevant cell type. The data were used to generate predictive models of the bioactive ligand environment within the brain interface neighborhood. Each box includes a ligand known to bind to a corresponding receptor expressed by at least one of the brain interface cell types. (A and B) show all of the ligands made by the brain interface cells that have a receptor expressed by at least one of the interface cell types. In (A) these are color-coded to show the cells that express these ligands while in (B) they are color-coded to show the cells that express the relevant receptor and thus are predicted to respond to the ligands. (C and D) show models with ligands expressed by the leptomeningeal cells that have corresponding receptors on either border macrophages (C) or border astrocytes (D). **(E)** Dot plot showing expression levels of *Igf2*, *Sema3a*, *Bmp6* and *Bmp7* mRNAs in leptomeningeal cells, border astrocytes and border macrophages, as determined from the scRNA-seq data. The size of the dot indicates the percentage of cells detectably expressing the mRNA, and the color indicates relative expression level, coded as per the adjacent keys. **(F)** Spatial plots of the leptomeningeal mRNAs *Bmp7, Bmp6, Igf2,* and *Sema3a* in all brain interface and layer one cells analyzed used the brain probeset (BP), shown on the same representative section. Relative mRNA expression levels are coded as per the adjacent keys. **(G)** High resolution Xenium Explorer images of the cortical interface region analyzed with the brain probeset (BP) showing expression of *Bmp6* (light pink dots), *Igf2* (blue dots)*, Bmp7* (green dots) and *Sema3a* (white dots) mRNAs relative to the leptomeningeal cells (pink), border astrocytes (yellow) and border macrophages (dark blue). The same field of view is shown in each panel, and arrows denote the same leptomeningeal cell detectably coexpressing multiple ligand mRNAs. Nuclei of other cell types (white) are also shown. Scale bars = 10um.

We also more specifically modeled ligands made by leptomeningeal cells that could influence their neighboring border astrocytes and macrophages (Fig. 7C, D; Extended Table 8). Most of the leptomeningeal ligands were predicted to act on both neighboring cell types. Analysis of the scRNA-seq data showed that some of these ligand mRNAs were highly enriched in the leptomeningeal cells relative to other neural and nonneural cell types, including *Igf2, Bmp6, Bmp7* and *Sema3a* (Fig. 7E; Fig. 7-1A). Since probes for these same ligands were present in the Xenium panels, we analyzed their spatial expression. Spatial plots showed that all four ligands were highly enriched and coexpressed in leptomeningeal cells at the brain interface (Fig. 7F, G). There was also scattered expression of these ligands in some layer one cells (Fig. 7F), consistent with the scRNA-seq. Notably, all of these ligands have been shown to directly affect astrocytes and macrophages (Aluganti Narasimhulu and Singla, 2020; Choe et al., 2014; Ishikawa et al., 1995) and cultured meningeal cells secrete all of these ligands (Choe et al., 2012; Ishikawa et al., 1995; Niclou et al., 2003; Ohe et al., 1996; Segklia et al., 2012). Thus, leptomeningeal cells express ligands that are predicted to directly influence their macrophage and border astrocyte neighbors and perhaps play a role in regulating the brain interface.

## Discussion

The meninges have been studied for many years, but we are only now gaining a cellular/molecular picture of this key interface tissue, largely due to the advent of high-resolution imaging approaches and single cell transcriptomics (Betsholtz et al., 2024). Most studies using these newer approaches have focused on the meninges and their vascular and immune cell components (Castro Dias et al., 2019; Como et al., 2023; DeSisto et al., 2020; Engelhardt et al., 2017; Pietilä et al., 2023; Rustenhoven et al., 2021). Here we have instead focused on the interface between the leptomeninges and the adjacent neural tissue. We show that in the adult cortex this interface is comprised of a layer of border astrocytes adjacent to leptomeningeal mesenchymal cells that are intermingled with resident macrophages and the occasional penetrating blood vessels. On the CNS side these interface cells are not closely-associated with other neural cells, although they may be in proximity to axons of passage in cortical layer one. This brain interface environment is predicted to be rich in growth factors, including ligands known to regulate interface cell types in other contexts. Thus, our data define and provide a molecular/cellular resource for the cortical interface, a unique brain compartment comprised of neural and nonneural cells that together regulate CNS interactions with the periphery (Choe et al., 2012; Chou et al., 2018; Como et al., 2023; DeSisto et al., 2020; Jordão et al., 2019; Lynn et al., 2011; Mapunda et al., 2022; Suter et al., 2017; Utz et al., 2020).

Data presented here confirm recent findings (DeSisto et al., 2020; Pietilä et al., 2023; Smyth et al., 2024) that the leptomeninges and dura are comprised of distinct types of mesenchymal cells. The dural cells that line the skull have much in common with connective tissue stromal cells and periosteal mesenchymal cells in other tissues. By contrast, the leptomeningeal cells are transcriptionally distinct from other mesenchymal cells and are differentially-enriched for tight junction (*Tjp1, Cdh5*) and gap junction (*Gjb2, Gjb6*) mRNAs, reflective of their unique biological function (Betsholtz et al., 2024; Castro Dias et al., 2019; Derk et al., 2022). We find that in the adult cortex the pial and arachnoid cells that comprise the leptomeninges are only a few cell layers thick, and that they are transcriptionally more similar to each other than they are to dural cells. We are, nonetheless able to distinguish them by expression of mRNAs like *Dpp4* and *Lama1*, and to confirm previous reports that pial but not arachnoid cells line the cortical hemispheric midline and are present on penetrating blood vessels (Bonney et al., 2022; Daneman and Prat, 2015; DeSisto et al., 2020; Jones et al., 2023; Pietilä et al., 2023).

Our data also define an intriguing population of astrocytes that interact with leptomeningeal cells at the cortical interface, in agreement with previous findings (Liu et al., 2013; Mason et al., 2021). These border astrocytes were enriched for 455 genes relative to cortical parenchymal astrocytes, including *Myoc* and *Gfap*. This unique astrocyte:pial cell relationship is reminiscent of astrocyte:vasculature cell interactions since in both cases astrocytic endfeet interact with an extracellular matrix layer deposited by adjacent nonneural cells (Daneman and Prat, 2015; Kadry et al., 2020; Rua and McGavern, 2018). However, while the vasculature-associated astrocytes are implicated in formation and maintenance of the blood:brain barrier, the function of brain interface border astrocytes is largely unknown (Batiuk et al., 2020; Mason et al., 2021). One likely function is to partner with pial cells and provide a cellular border that ensures cells do not move between neural and non-neural compartments, analogous to an epithelium. However, developmental studies suggest that border astrocytes may be more than just structural. During embryogenesis the endfeet of cortical radial glial precursors form a similar interface with the leptomeningeal cells and this interplay is essential for appropriate cortical development (Chou et al., 2018; Halfter et al., 2002; Myshrall et al., 2012).

One somewhat surprising finding reported here is that border astrocytes are the only CNS cell type in close proximity to leptomeningeal cells. One potential explanation for this finding is that astrocytic endfeet spatially exclude other cell types from the interface. However, there may also be a more active exclusion mechanism. For example, we show that the known repulsive ligand Sema3b is highly enriched in border versus parenchymal astrocytes, and this molecule could serve to repel axons and other neural cell types from the interface region (Ducuing et al., 2020; Nawabi et al., 2010).

The third major cell type within the brain interface neighborhood is an intriguing population of leptomeninges-resident macrophages. As previously reported (Jordão et al., 2019; Sankowski et al., 2024) we find that these are transcriptionally-distinct from both the cortical dural macrophages (data not shown) and from microglia within the cortical parenchyma. The spatial transcriptomic data show that these resident macrophages are embedded within the leptomeningeal layers, in close proximity to both leptomeningeal cells and border astrocytes. Given their location, these macrophages are ideally situated to play a role in barrier function, and to act as a first-line response to injury (Jordao et al 2019; Joost et al. 2019; Pedragosa et al 2018). Our ligand-receptor modeling suggests they may also directly regulate leptomeningeal cell and border astrocyte biology via secretion of ligands like TNF and Il1beta. Since these ligands are known to regulate the biology of mesenchymal cells in other tissues, then their relative importance for interface biology will be an important future avenue of investigation (Magliozzi et al., 2019; Mokbel et al., 2024).

Our spatial transcriptomic data also provide a cellular level view of layer one cortex, and define microglia, OPCs, parenchymal astrocytes, and Lamp5-positive interneurons as the major cellular residents. On the surface side layer one is bordered by brain interface astrocytes and on the other side by relatively dense layer 2 excitatory neurons, interneurons and oligodendrocytes, consistent with previous morphological studies (Yao et al., 2021). Notably, we find that layer one cells are largely not in close proximity to each other. Since layer one includes many axons of passage, then it is possible that axonal interactions comprise the major homeostatic association for most layer one cells. Moreover, the relative distance between border astrocytes and other layer one cells suggests that secreted ligands may be the major way the brain interface communicates with the underlying cortical parenchyma (Gesuita and Karayannis, 2021; Suter et al., 2017).

Our findings indicate that the brain interface is a unique cellular neighborhood comprised of three major cell types, leptomeningeal cells, border astrocytes and tissue-resident macrophages. It is likely that communication between these closely-associated cellular players involves both cell:cell contacts and secreted ligands. Our modeling and spatial transcriptomics provide growth factor candidates for this communication. As one example, we show that *Bmp6* and *Bmp7* expression are highly-specific to adult leptomeningeal cells, and others have shown that these ligands are secreted by cultured leptomeningeal cells (Niclou et al., 2003) and can regulate both astrocyte and macrophage biology (Aluganti Narasimhulu and Singla, 2020; Choe et al., 2012; Du et al., 2019; Sabo et al., 2009). As a second example, we show that *Sema3a* is specifically expressed by adult leptomeningeal cells, and others have shown that meningeal cells synthesize Sema3a protein (Niclou et al., 2003) and that it is anti-inflammatory for macrophages (Rienks et al., 2017) and regulates glial precursor biology (Sugimoto et al., 2001). Thus, our predictive cell interaction and growth factor models, together with the detailed cellular-molecular definition of interface cells and their locations, will provide the basis for future studies asking how interactions between these three distinct brain interface cell types allow them to maintain the specialized brain:periphery interface, keep cells from transitioning in and out of the brain, and repair/regenerate after brain injury.

## Author Contributions

S.N.E. designed research, performed research, analyzed data and wrote the paper; C.E. designed and performed research; K.K. and L-P.B. performed research; D.R.K. and B.A.M. designed research; F.D.M. designed research, analyzed data and wrote the paper.

## Acknowledgements

We thank Ashleigh Willis and Jasmine Yang for technical advice and experimental help.

## Conflict of interest statement

The authors declare no conflict of interest.

## Funding sources

This work was funded by CIHR grants to F.D.M., D.R.K. and B.A.M. S.E. was funded by a CGSM studentship.

## Extended Figure Legends

**Extended Figure 1-1.**
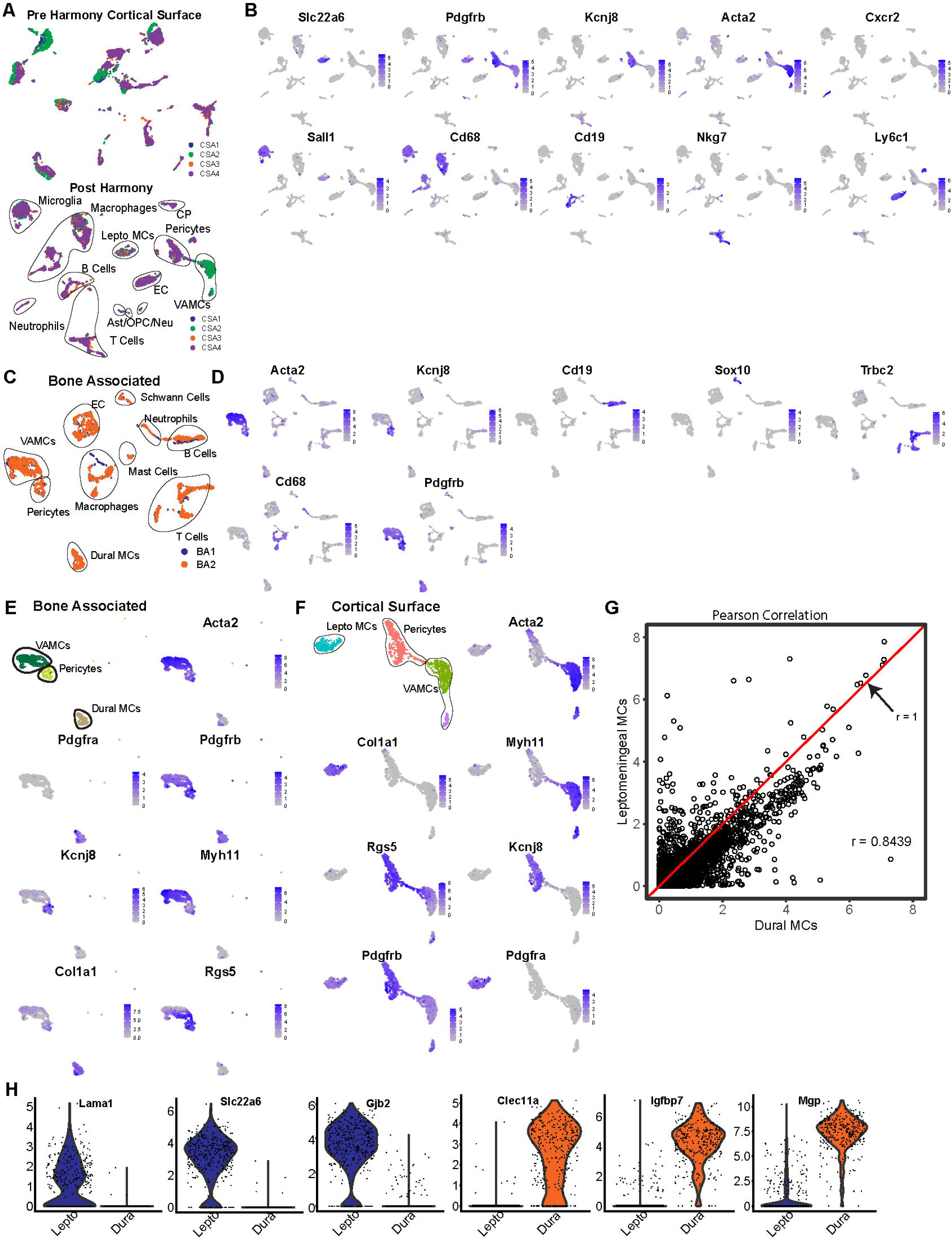
scRNA-seq to analyze adult murine cortical leptomeninges and dura-associated cell types. **(A)** UMAP visualization of merged transcriptomes from 4 independent cortical surface-associated (CSA) scRNA-seq runs (CSA1-4). Transcriptomes from each run are colored as per the adjacent legend. The top panel shows the dataset without Harmony batch correction, and the bottom with one iteration of batch correction. The bottom UMAP is also annotated for cell types. **(B)** The batch-corrected, merged dataset in (A, bottom panel) was overlaid for expression of marker genes specific to different cell types. Expression levels are coded as per the adjacent keys. **(C)** UMAP visualization of merged transcriptomes from 2 independent bone-associated scRNA-seq runs (BA1, BA2). Transcriptomes from each run are colored as per the adjacent legend. The UMAP is also annotated for cell types. **(D)** The merged dataset in (C) was overlaid for expression of marker genes specific to different cell types. Expression levels are coded as per the adjacent keys. **(E, F)** *Pdgfrb*-positive mesenchymal cells were subsetted from the bone-asociated and cortical surface-associated datasets as shown in (A) and (C), and reanalyzed. The annotated UMAPs in the upper left corners show the subsetted transcriptomes colored by cluster, and annotated for *Pdgfra*-positive dural or leptomeningeal mesenchymal cells (Dural Mes in E, Lepto Mes in F), pericytes or vasculature-associated *Pdgfra*-negative mesenchymal cells (VAMCs). The remainder of the panels show expression overlays for genes characteristic of each of the different mesenchymal cell types. Expression levels are coded as per the adjacent keys. **(G)** Pearson correlation analysis of averaged expression of each detected gene in leptomeningeal (y-axis) versus dural (x-axis) mesenchymal cell transcriptomes from the merged dataset shown in (E and F). **(H)** Violin plots showing relative expression levels of select mRNAs that were identified in the differential gene expression analysis as being highly-enriched in leptomeningeal (colored blue) versus dural (colored orange) cells, respectively.

**Extended Figure 2-1.**
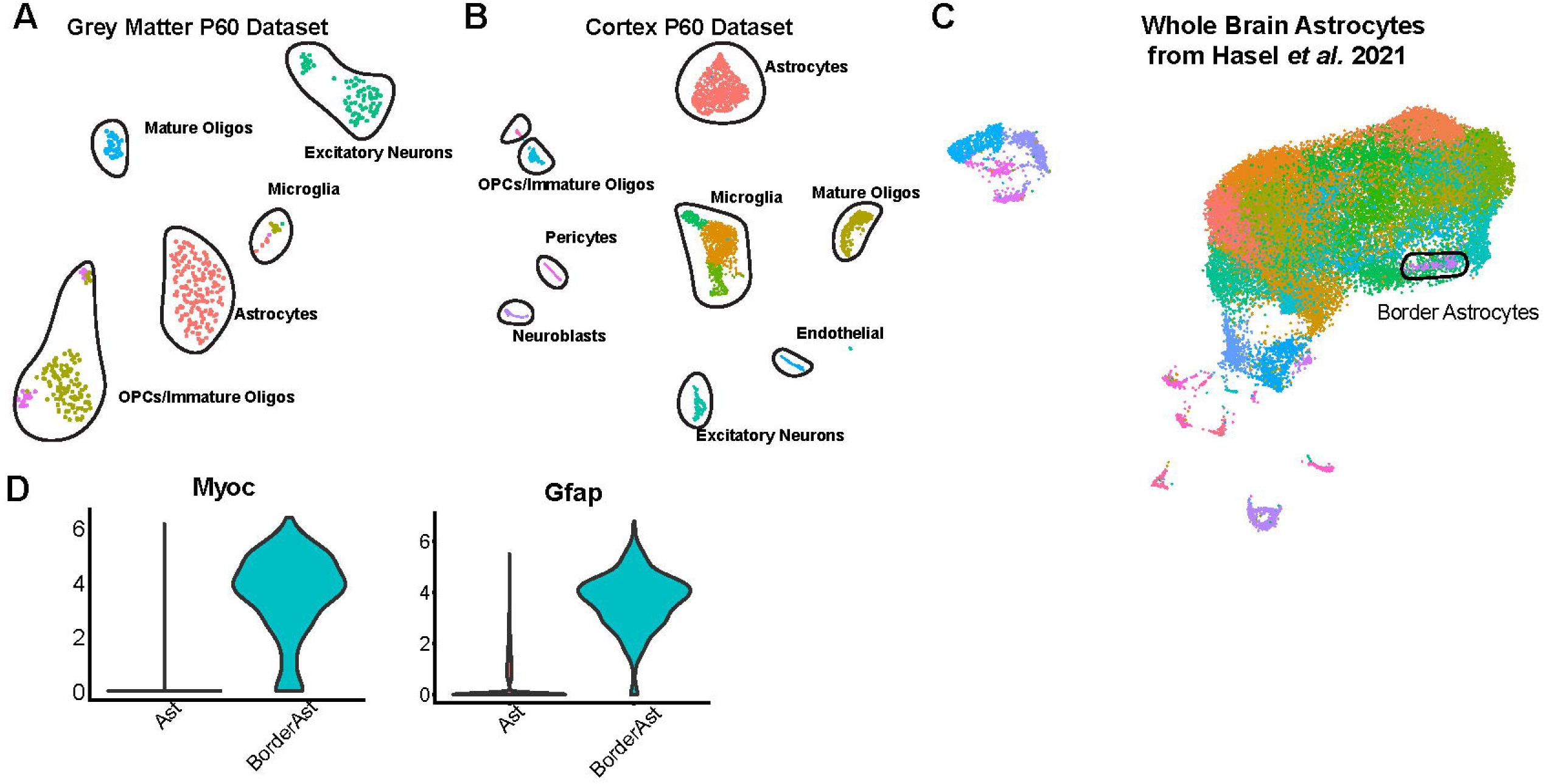
scRNA-seq to analyze adult cortical and border astrocytes. **(A, B)** UMAP visualizations of transcriptomes from previously-published (Daniel et al., 2024; GEO GSE255405) scRNA-seq datasets of dissected postnatal day 60 (P60) cortical grey matter (A) or total cortex (B) tissue. Transcriptomes were reanalyzed and cell types were identified using well-characterized marker genes. The transcriptionally-distinct clusters are color-coded, and different cell types annotated and denoted by the solid lines. **(C)** UMAP visualization of transcriptomes from a previously-published (Hasel et al., 2021; GEO GSE148611) whole brain astrocyte scRNA-seq dataset. Transcriptomes were reanalyzed and transcriptionally-distinct clusters are color-coded. The cluster most enriched for *Myoc* and *Gfap* expression (outlined) was subsetted and used for subsequent analyses. **(D)** Violin plots showing relative expression levels of *Myoc* and *Gfap* mRNAs in the subsetted putative whole brain border astrocytes (colored in blue; as outlined in C) versus all other astrocytes in (C) (colored in red).

**Extended Figure 3-1.**
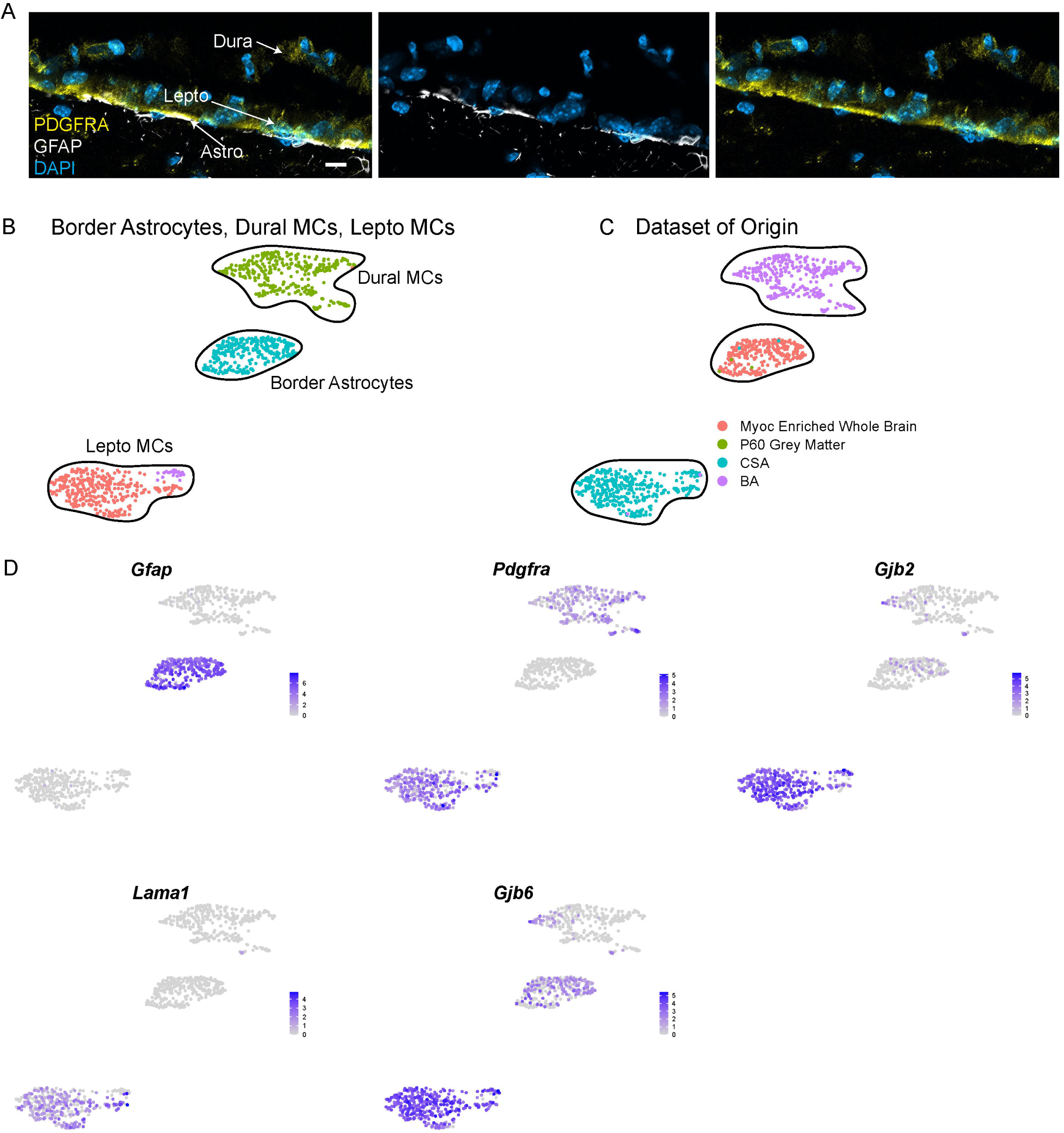
Characterization of the cortical brain interface by immunostaining. **(A)** High magnification confocal images of a coronal section through the brain interface and skull, immunostained for PDGFRα (yellow) and GFAP (white), and counterstained with DAPI (blue). The left panel shows the merge and the right two immunostaining for PDGFRα or GFAP separately, as indicated. Lepto = leptomeninges, Astro = border astrocytes. Scale bar = 10um. **(B, C)** The border astrocyte transcriptomes from cluster 2 in Figure 2A and the leptomeningeal and dural mesenchymal cell transcriptomes from Figure 1E were merged and reanalyzed. Shown are UMAPs with the clusters color-coded and annotated (B) or the datasets of origin indicated as per the adjacent color legend (C). CSA = cortical surface-associated dataset, BA = bone-associated dataset. **(D)** Gene expression overlays of the merged astrocyte plus meningeal mesenchymal cell dataset shown in (B, C) for selected genes that distinguish the different cell types. Expression levels are color-coded as per the adjacent keys.

**Extended Figure 4-1.**
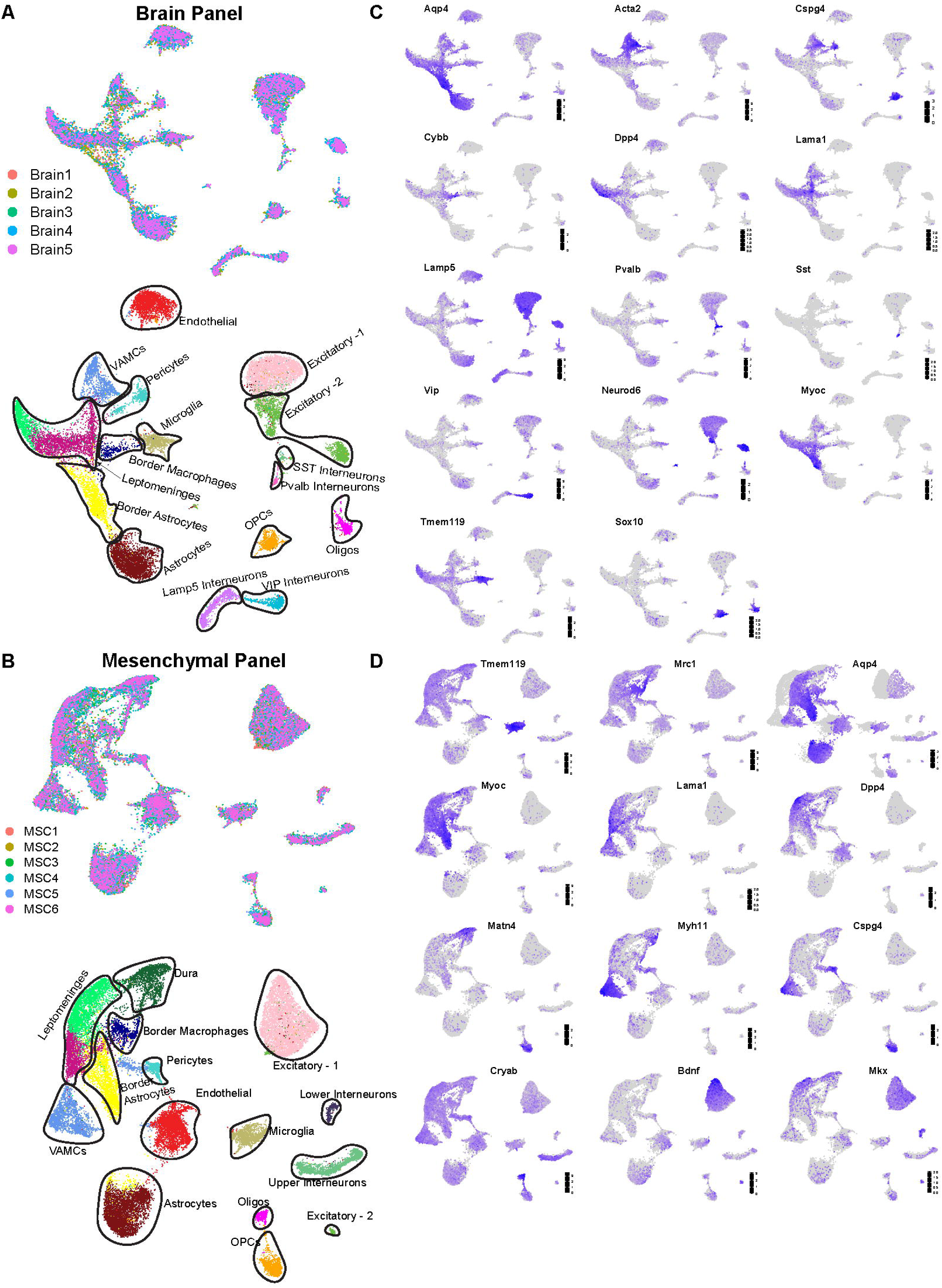
Analysis of the cortical interface and layer one cells using single cell multiplexed in situ gene expression analysis with the brain and mesenchymal probesets. Coronal cortical sections were analyzed by Xenium-based single cell multiplexed *in situ* gene expression analysis with either brain-targeted or mesenchymal cell-targeted probesets. The ROI that was analyzed is shown in Figure 4A, and the annotated UMAP cluster visualization of the resultant merged transcriptomes shown in Figure 4B and C. **(A, B)** UMAPs showing the merged brain probeset (A) or mesenchymal probeset (B) datasets showing either the annotated clusters (bottom; the same UMAPs as in Figure 4B and C) or the section of origin (top) for each of the transcriptomes (right). Each dot represents a single transcriptome. VAMC = vasculature-associated mesenchymal cell, Oligos = oligodendrocytes, SST = somatostatin, Pvalb = parvalbumin, VIP = vasoactive intestinal peptide. Brain1-5 indicates 5 sections from 5 different mice. MSC1-6 indicates 6 sections from 4 different mice. **(C, D)** Expression overlays for selected marker genes on the UMAPs shown in (A, B). Expression levels are coded as per the adjacent keys.

**Extended Figure 6-1.**
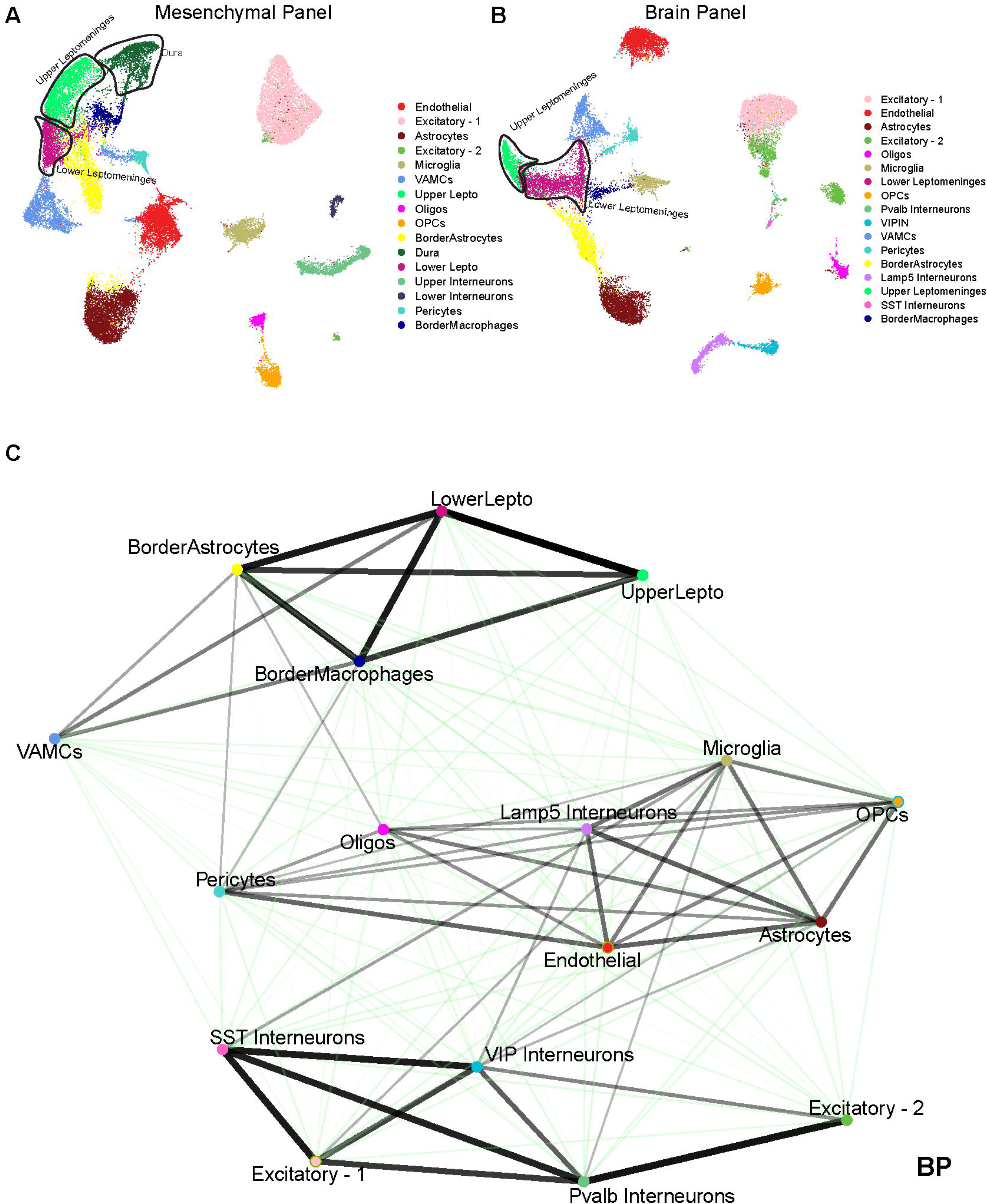
Defining the brain interface and layer one cellular neighborhoods using single cell spatial transcriptomics. **(A, B)** UMAPs showing the single cell spatial transcriptomic datasets from Figure 4B and C analyzed at higher resolution (0.8 for the Brain Probeset and 1.2 for the Mesenchymal Probeset) to show separate clustering of upper versus lower leptomeningeal cell transcriptomes. Shown are merged transcriptomes from analyses with the mesenchymal cell (A) or brain (B) probesets. Transcriptionally-distinct clusters are color-coded and annotated for cell types as per the adjacent legends. **(C)** Proximity analysis showing the relative strength of interactions between different cell types located in the cortical brain interface and adjacent layer one, as analyzed from the dataset obtained using the brain probeset. The nodes are color-coded and annotated for cell type, and the weight of the line indicates the strength of the interaction. Black lines indicate significant interactions based on a permutation test, comparing our data to a null distribution created from the random permutation of cell labels with fixed positions (See Methods and Materials for details). Green lines indicate interactions that were not statistically significant. Excitatory neuron-1 and excitatory neuron-2 denote transcriptionally-distinct excitatory neuron clusters. Cell centroid information was used to perform this analysis, with a proximal interaction being defined as one where cell centroids were within 70µm of another cell centroid. Oligos = oligodendrocytes, Pvalb = parvalbumin, SST = somatostatin, Lepto = leptomeningeal cells, VIP = vasoactive intestinal peptide.

**Extended Figure 7-1.**
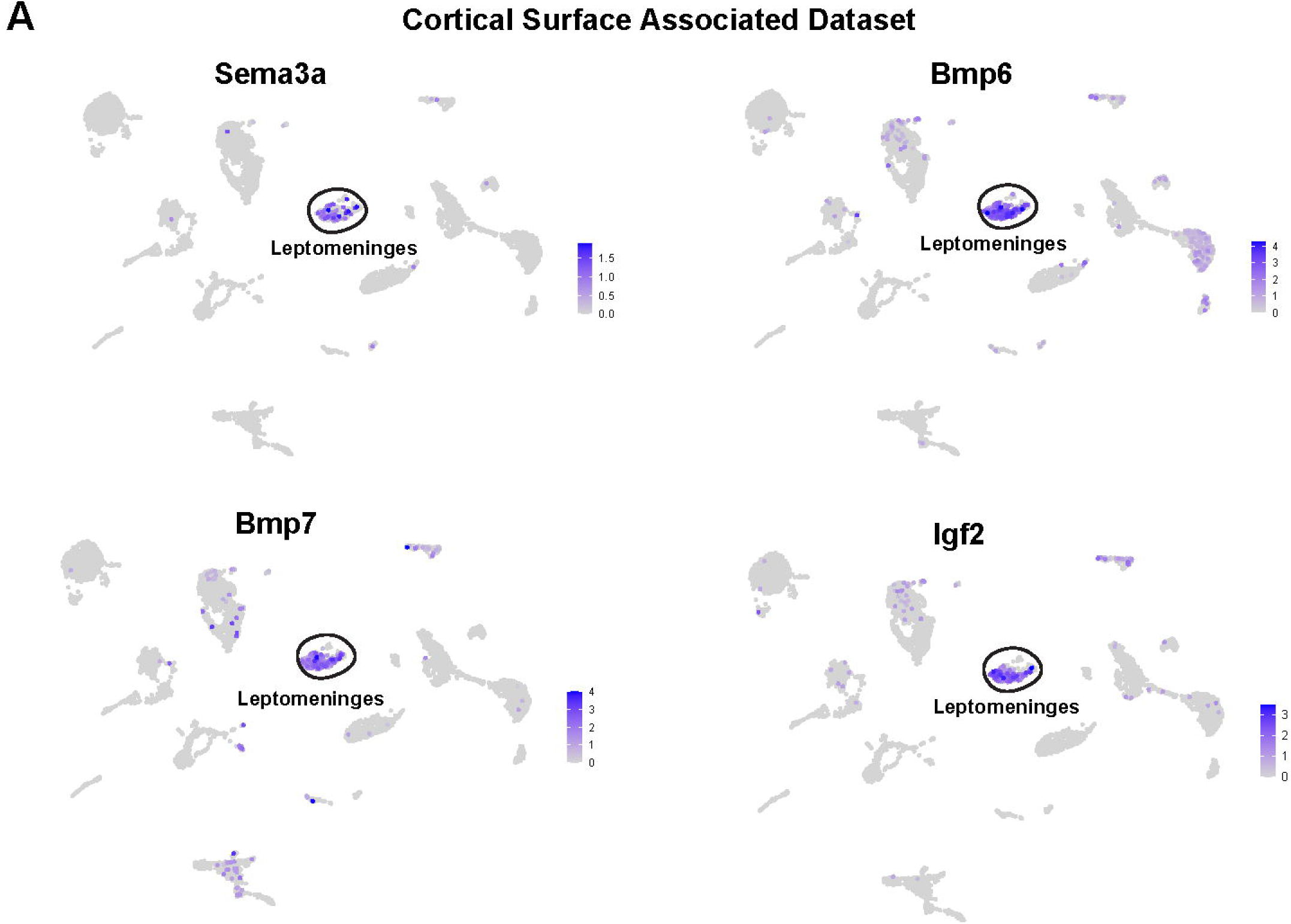
Specific expression of leptomeningeal ligands, as analyzed by scRNA-seq. **(A)** UMAPs showing the cortical surface-associated dataset from Figure 1A, overlaid for expression of four ligands that are highly-enriched in the leptomeningeal cells, *Sema3a, Bmp6, Bmp7* and *Igf2*. Expression levels are color-coded as per the adjacent keys.

**Extended Table 1. *Genes differentially expressed in leptomeningeal and dural mesenchymal cells (MCs).*** Differential gene expression analysis was performed on the merged Pdgfra+ clusters in the Cortical Surface Associated and Bone Associated Datasets as shown in Figure 1E. Differentially expressed gene lists were defined as those expressed in >10% of cells in either group with a Bonferroni adjusted p-value < 0.05 (Adj. p-value) and ≥ 1.3 average log2 fold change (avg log2FC).) * Denotes genes included in the leptomeningeal mesenchymal cell signature. ** Denotes genes included in the dural mesenchymal cell signature.

**Extended Table 2. *Gene ontology for genes differentially enriched in leptomeningeal and dural mesenchymal cells as shown in Extended Table 1.*** g:Profiler was used to determine what gene ontology terms were overrepresented in the transcripts differentially expressed between leptomeningeal and dural mesenchymal cells. Enriched terms from the Molecular Function (MF), Biological Process (BP), and Cell Components (CC) categories are all included.

**Extended Table 3. *Genes differentially expressed in border astrocytes versus cortical grey matter astrocytes.*** Differential gene expression analysis was performed on the merged astrocyte dataset shown in Figure 2A, comparing the border astrocytes (cluster 2) and the cortical grey matter astrocytes (cluster 1). Differentially expressed gene lists were defined as those expressed in >10% of cells in either group with a Bonferroni adjusted p-value < 0.05 (Adj. p-value) and ≥ 1.3 average log2 fold change (Avg log2FC). *Denotes genes included in the border astrocyte gene signature.

**Extended Table 4. *Gene ontology for genes differentially enriched in border astrocytes versus cortical grey matter astrocytes as shown in Extended Table 3.*** g:Profiler was used to determine what gene ontology terms were overrepresented in the transcripts differentially expressed between border astrocytes and cortical grey matter astrocytes. Enriched terms from the Molecular Function (MF), Biological Process (BP), and Cell Components (CC) categories are all included.

**Extended Table 5. *Genes differentially expressed in border macrophages versus cortical microglia.*** Differential gene expression analysis was performed on the border macrophages and microglia from the Cortical Surface Associated and Bone Associated Datasets as shown in Figure 1A and B. Differentially expressed gene lists were defined as those expressed in >10% of cells in either group with a Bonferroni adjusted p-value < 0.05 (Adj. p-value) and ≥ 1.3 average log2 fold change (Avg log2FC).

**Extended Table 6. *Analyses of leptomeningeal cell, border macrophage and border astrocyte ligand mRNA expression.*** Shown are ligand mRNAs expressed in leptomeningeal mesenchymal cells, border macrophages and border astrocytes at the cortical interface as determined by scRNA-seq, as well as the percentage of a given cell type that detectably expresses that ligand. For leptomeningeal cells and border macrophages ligands were extracted from the cortical surface-associated and bone-associated scRNA-seq datasets shown in Figure 1A and B, and for border astrocytes from the merged astrocyte scRNA-seq dataset shown in Figure 2A based on a curated ligand and associated receptor database (Toma et al., 2020). Ligands were only considered for further analysis (such as in the ligand-receptor modeling shown in Extended Table 8) if they were detectably expressed in at least 5% of the relevant cell type.

**Extended Table 7. *Analyses of leptomeningeal cell, border macrophage and border astrocyte receptor mRNA expression.*** Shown are receptor mRNAs expressed in leptomeningeal mesenchymal cells, border macrophages and border astrocytes at the cortical interface as determined by scRNA-seq, as well as the percentage of a given cell type that detectably expresses that ligand. For leptomeningeal cells and border macrophages receptor mRNAs were extracted from the cortical surface-associated and bone-associated scRNA-seq datasets shown in Figure 1A **and** B, and for border astrocytes from the merged astrocyte scRNA-seq dataset shown in Figure 2A based on a curated ligand and associated receptor database (Toma et al., 2020). Receptors were only considered for further analysis (such as in the ligand-receptor modeling shown in Extended Table 8) if they were detectably expressed in at least 5% of the relevant cell type.

**Extended Table 8. *Details of ligand-receptor communication models.*** Predicted ligand and receptor communication models were generated using the leptomeningeal, border macrophage and border astrocyte ligand and receptor mRNA lists shown in Extended Tables 6 and 7. Shown are ligand-receptor communication models of (8-1) ligands from all cells predicted to act on leptomeningeal, border macrophage or border astrocyte receptors, (8-2) leptomeningeal ligands predicted to act on border macrophage receptors, and (8-3) leptomeningeal ligands predicted to act on border astrocyte receptors.

**Extended Table 9. *Xenium probesets.*** Shown are the mesenchymal custom probeset, comprised of probes for 480 genes, as well as the custom add-on brain probeset genes that were used in conjunction with the 10X predesigned 247 probe mouse brain panel. The numbers of probes per gene were tuned for each Xenium custom add-on panel to ensure robust detection and isoform coverage, while avoiding optical crowding. Probe number selection was informed by the 10X Xenium panel designer.

## Notes

### Competing Interest Statement

The authors have declared no competing interest.

